# RNA–X: Modeling RNA interactions to design binder RNA and simultaneously target multiple molecules of different types

**DOI:** 10.1101/2025.11.24.690191

**Authors:** Sobhan Shukueian Tabrizi, Helyasadat Hashemi Aghdam, Kerim Eren, Cansu Yilmaz, Duygu Kuzuoglu-Ozturk, A. Ercument Cicek

**Affiliations:** Department of Computer Engineering, Bilkent University, Ankara, Türkiye; Faculty of Engineering and Natural Sciences, Sabancı University, İstanbul, Türkiye; Department of Urology, University of California, San Francisco, San Francisco, CA, USA; Helen Diller Family Comprehensive Cancer Ctr., University of California, San Francisco, San Francisco, CA, USA

## Abstract

RNA interactions with proteins, other RNA molecules, and DNA play essential roles in numerous cellular processes and underpin a wide range of therapeutic mechanisms. Consequently, modeling these interactions is critical for understanding biological systems and designing novel RNA-based therapeutics. However, designing RNA sequences that selectively and strongly bind to a specific target remains a major challenge due to the vast sequence space and the current limitations of experimental and computational methods. In this study, we introduce RNA–X, the first RNA interaction foundation model which is based on masked language modeling for representation learning, conditional RNA design, and optimization. For the first time, RNA–X enables (i) RNA design targeting not only proteins but also other RNA and DNA molecules, and (ii) simultaneous design against multiple targets. Our extensive experiments demonstrate that RNA–X generates RNA sequences with natural structural characteristics and surpasses state-of-the-art methods in protein targeting. We experimentally validate RNA–X in vitro, where a RNA–X-designed RNA targeting the 5′UTR of the androgen receptor (*AR*) mRNA was shown to reduce both *AR* transcript and protein levels in prostate cancer cells. Using affinity predictors and molecular dynamics simulations, we further show that the model can design RNA molecules that (i) target therapeutically relevant molecules such as p53 and thrombin proteins, and (ii) bind to targets with no prior interaction data, such as PHF6 protein. As a proof of concept, we designed a novel guide RNA from scratch that simultaneously binds (i) to the DNA of a bacterium and to the Cas9 protein. The resulting design achieves a predicted binding energy comparable to that of the wild-type guide RNA reported in the Protein Data Bank. Despite having orders of magnitude fewer parameters than existing RNA foundation models, RNA–X produces representations that outperform them across diverse RNA interaction–related downstream tasks. The code and pretrained model are publicly available on GitHub.

## 1 Introduction

RNA interactions play a central role in various biological systems such as (i) microRNA–mRNA interactions for post-transcriptional regulation [4], (ii) long noncoding RNA (lncRNA) - DNA interactions for the formation of molecular scaffolds to change chromatin structure [35], and (iii) RNA - protein interactions to regulate splicing [44] and transport [45] [16]. Consequently, the design of novel RNA sequences that can interact with target molecules presents ample opportunities in therapeutic applications.

The therapeutic potential of RNA design through binding to a target X (hence the name RNA-X) has many examples for various molecule types. **RNA-RNA**. In RNA interference (RNAi), small interfering RNAs (siRNAs) silence disease-related genes through degradation of their precursor mRNA. Yet, the design of potent siRNAs remains challenging due to requirements for high efficacy, low off-target effects, and minimal immune activation [1]. **RNA-DNA**. CRISPR–Cas systems rely on single-guide RNAs (sgRNAs) to direct Cas proteins to genomic loci. Ensuring sgRNA efficacy and specificity in design is crucial and challenging to ensure reliable gene editing. **RNA-protein**. RNA binding proteins (RBPs) offer a strategy to inhibit or enhance RNA–protein interactions. This approach expands the current drug landscape, as only a small fraction of human proteins are targetable using ligand-based drugs[20]. Despite its potential, designing RNA molecules that selectively bind to different types of targets remains an open challenge, mostly because of the vast search space and stringent efficacy and specificity requirements. Current practice in the lab relies heavily on trial and error such as high-throughput screening techniques such as SELEX (Systematic Evolution of Ligands by EXponential Enrichment) [5]. While computational methods can help designing RNA to bind with other molecules process, they have critical limitations. Recent computational research on RNA is focused on the building of RNA foundation models [6,33,50,53,13].

However, most of these models are not generative and cannot directly design new sequences conditioned on a target [40]. Early generative methods focused on inverse folding, where sequences are designed to fold into a desired secondary structure [19,15,2,18,38,11,8]. Various new deep learning-based models have been proposed to design the RNA structure or sequence that resembles natural RNA [14,36,43,24,3,21,49,23,7]. However, these methods cannot condition the design to bind with a target. RNAFlow is a diffusion-based model that can design RNA structure conditioned on a target protein [30]. RNAGen [31] and GenerRNA

[52] are earlier examples of RNA sequence designers to target a protein which condition for the target only in post-processing, using affinity predictors to guide generation or fine-tuning the model for the target using interaction data. This creates key limitations: (i) the generation and optimization are decoupled, preventing joint RNA–target representations; (ii) target-specific predictors have a narrow coverage and for many proteins there are no known RNA-interactions: For example, DeepCLIP [17] supports only 221 proteins; and the design of RNAs for new or synthetic targets is infeasible without existing interaction data. Recent approaches focus on directly modeling RNA-protein interactions to avoid these problems. BAnG uses protein structure and sequence as input and trains a model on 3D RNA-protein complex structure data in the Protein Databank (PDB) [26]. RNAtranslator is a language model that frames the conditional RNA design problem as a sequence-to-sequence natural language translation problem [41]. InstructNA couples nucleic-acid language models with high-throughput SELEX data to design aptamers de novo [39], but remains protein-only and requires target-specific selection data. Despite progress, all of the methods mentioned above can design RNA to bind to a single target type (proteins) and their embeddings cannot be used for downstream RNA interaction-related tasks, as they are not designed as foundation models for representation learning.

In this study, we present RNA–X, the first RNA interaction foundation model. RNA–X can design an RNA sequence to target RNA, DNA, and proteins, and can design a single RNA molecule to simultaneously bind to multiple targets for the first time. RNA–X, is a masked language model trained end-to-end on more than hundred million RNA-target interactions. It learns joint representations of RNA and target molecules, enabling direct sampling of RNA sequences with desired binding properties. We formulate the problem of target-conditional RNA design as a context-aware generative task based on a target-specific embedding. RNA sequences are generated using iterative unmasking and refinement of the tokens coupled with a Monte Carlo tree search informed by model’s rewards. The model representations are further optimized to handle the representation degeneration in sequence level representations using whitening and uniformity for increased performance in downstream tasks without changin model weights. See Figure1 for RNA–X system model. Our results show that RNA–X designs natural-like RNAs (i) with matching or better predicted affinity/stability compared to known binders of all target types. It matches or outperforms the state-of-the-art methods across seventeen protein targets, four of which are excluded entirely from its training set. We further design binding RNAs for 71 additional proteins, where RNA–X generates higher predicted affinity than random sequences for 50 of them (Supplementary Figures 11–12). We experimentally validate the RNA–RNA design setting in human prostate cancer cells, where a RNA–X-designed RNA targeting the *AR* 5′UTR downregulates the gene at both the transcript and the protein level. We show that the model can successfully design to target therapeutically relevant molecules such as p53 and thrombin proteins, and for targets without prior interaction data (e.g., PHF6 protein), enabling the design for virtually any target molecule sequence. For the first time, we designed a novel guide RNA from scratch in a single shot that simultaneously binds to (i) the DNA of Streptococcus pyogenes, and (ii) the Cas9 protein. The resulting design achieves a predicted binding energy comparable to that of the wild-type guide RNA reported in the Protein Data Bank and is close to a manual optimization. Despite having orders of magnitude fewer parameters than existing RNA foundation models, our optimized embeddings outperform the state-of-the-art in RNA interaction–related downstream tasks such as estimating siRNA efficacy and sgRNA on-target knockout efficacy prediction. The code and pretrained model are publicly available on GitHub.

## 2 Results

### 2.1 Overview of RNA–X

RNA–X is a masked language model trained end-to-end on more than one hundred million RNA-target interactions, as shown in Figure 1 (Architecture). It learns joint representations of RNA and target molecules, enabling direct sampling of RNA sequences with desired binding properties. Since RNA folding shows longrange dependencies, where local nucleotide changes can induce global structural rearrangements, RNA–X benefits from conditioning on both the left and right context of the RNA sequence in addition to the target sequence. Additionally, RNA sequence design is a discrete optimization problem in which objectives such as minimum free energy minimization or secondary structure properties are typically non-differentiable with respect to sequence. To address this challenge, we formulate target-conditional RNA design as a context-aware generative task based on target-specific embeddings and optimize candidate sequences using a Monte Carlo tree search (MCTS) procedure guided by RNA–X’s learned proposal distribution and scoring functions. This formulation enables position-wise editing during optimization, allowing the search process to evaluate the effect of single-nucleotide changes on the objective while permitting modifications at arbitrary sequence positions. As a result, the framework provides flexible exploration of the sequence landscape and supports fine-grained refinement of candidate RNAs. RNA sequences are generated through iterative token unmasking and refinement, coupled with model-informed search to efficiently explore the combinatorial sequence space (Figure 1, Inference). Unlike direct target-to-RNA translation approaches that generate sequences in a single pass, our formulation treats RNA design as an iterative optimization process, enabling non-local edits and explicit evaluation of nucleotide-level changes during search.

**Fig. 1:**
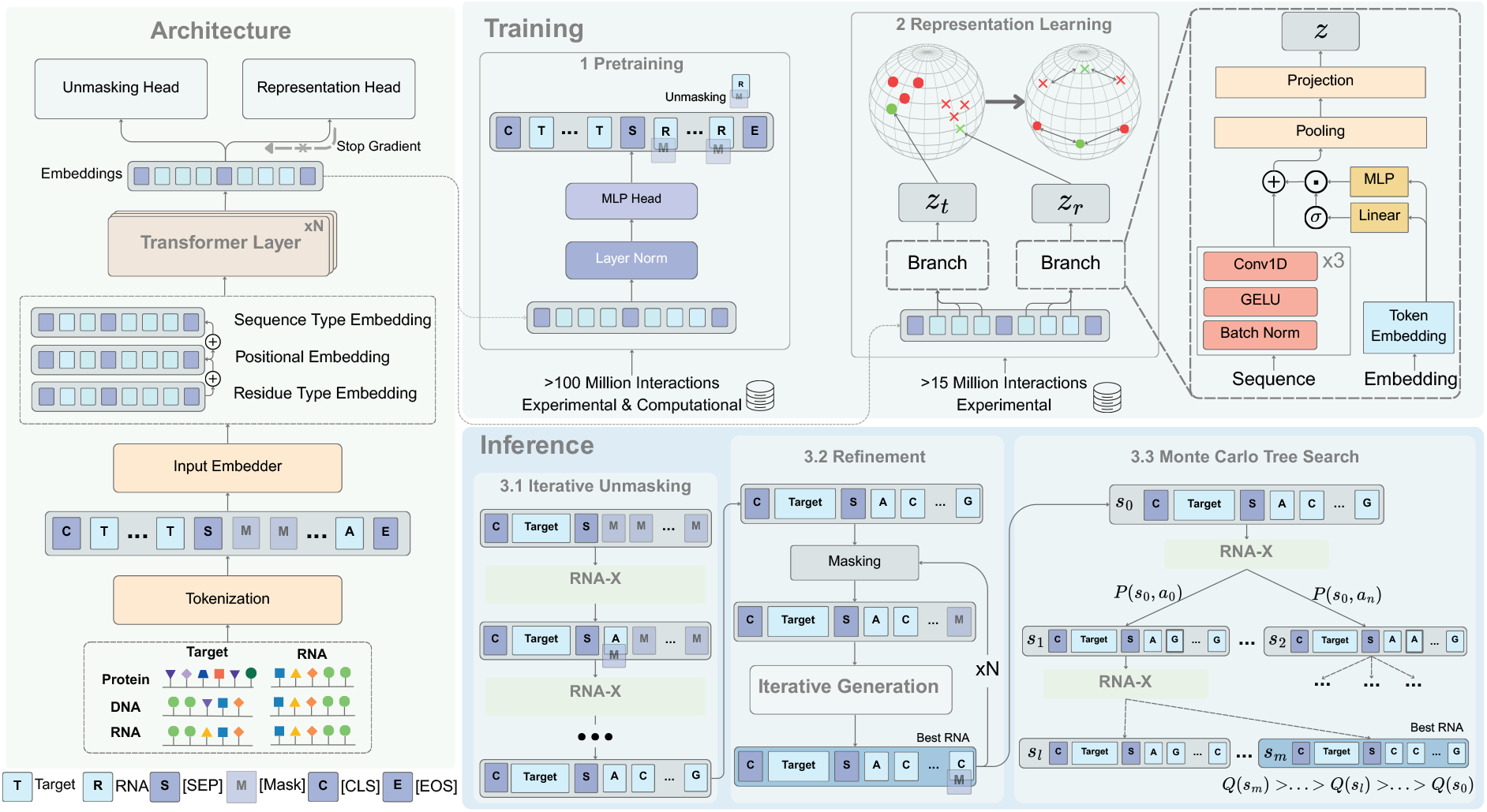
Overview of RNA–X. Architecture: RNA–X encodes paired RNA–target sequences through a shared transformer backbone with sequence-type, positional, and residue-type embeddings. Separate unmasking and representation heads enable both generation and representation learning objectives. **Training:** The model is trained on over 100 million RNA–target interactions. (1) RNA–X is pretrained by unmasking the masked tokens. Then (2) interaction-aware representation learning using contrastive branches for target (*z*_*t*_) and RNA (*z*_*r*_) is performed. The representations obtained in pretraining are transformed into another space (transformed globe) to better distinguish interacting pairs and have better downstream performance. Model weights are kept frozen at this stage. **Inference:** During generation, RNA–X takes target sequence(s) (protein, RNA, DNA, or combinations) and fully masked RNA sequence. It designs RNA sequence through iterative unmasking of the tokens from left to right (3.1). The result is repeatedly refined by randomly masking tokens and passing them through iterative unmasking (3.2). Finally, a Monte Carlo Tree Search (MCTS)–based exploration is performed (3.3). The probability *P* of choosing action *a* in state *s* is deter-mined by RNA–X logits, and the value of the state *Q* is calculated using the scoring function (Supplementary Notes 1.7), which optimizes the designed sequences to bind the target while preserving RNA structural and thermodynamic stability.

We further train the model to learn sequence-level representations by aggregating token embeddings learned by RNA–X, as shown in the Training panel of Figure 1, aiming to capture interaction-based molecular features while handling the representation degeneration problem. To improve the informativeness of these embeddings, we apply post hoc whitening and uniformity regularization, which enhances downstream performance without changing the underlying model weights. The learned sequence-level representations provide a unified interface for downstream interaction-related tasks, such as binding affinity prediction. Additionally, for tasks with sufficient labeled data, lightweight prediction modules can be trained on top of these representations and integrated into the optimization loop as scoring functions, enabling objective-specific sequence refinement during search. In this way, training a representation learning head on top of the frozen pretrained model requires less data and is more efficient than training a separate model, while also ensuring that both sequence generation and downstream evaluation (scoring) operate within the same embedding space.

### 2.2 RNA–X designs RNAs with higher predicted binding score compared to other generative models

We compare RNAs designed by RNA–X with RNAs designed by other models when targeting proteins. Please note that there are no other methods in the literature that can be compared when targeting RNA or DNA. For this comparison, we train a separate DeepCLIP model [17] for each target, using a dedicated training set and an independent test set to ensure unbiased predictions (Section 4.1). DeepCLIP’s reliability depends on the training data, and CLIP datasets can carry protocol- and batch-specific biases. Because combining datasets across protocols improves robustness [32], we pool CLIP data from multiple experiments for each target before splitting. We retain only evaluators reaching an AUROC of at least 0.80 on their test set and discard the rest, which is what determines the set of proteins we can evaluate. We then design 100 sequences per target with each method and score them with that target’s DeepCLIP model. Since the evaluators are trained and tested independently of the designed sequences, this evaluation remains unbiased. We benchmark against four recent models: GenerRNA, RNAGEN, BAnG and RNAtranslator. These differ in what they require of the target. GenerRNA must be fine-tuned on target-specific binding data, RNAGEN relies on an external DeepBind predictor to guide optimization, and BAnG needs the target structure, predicted with AlphaFold2 [25] when unavailable. RNAtranslator, like RNA–X, needs only the target sequence. We exclude RNAFlow, which the BAnG manuscript shows BAnG outperforms.

For a comprehensive comparison we consider three groups of proteins. First, we consider two RNA-binding proteins (RBPs) with substantially different numbers of known interactions in CLIPdb: RBM5, with a relatively small set of 3, 917 interactions, and ELAVL1, with a much larger set of 1, 079, 145. This lets us assess how the amount of available interaction data affects the designs. Because RNAGen and GenerRNA require per-target fine-tuning, which demands substantial computational resources, we restrict the benchmark with these two methods to RBM5 and ELAVL1. RNAtranslator and BAnG have no such restriction, so we benchmark against them on fifteen further proteins, selected among proteins with enough known interactions to train a reliable evaluator. Eleven of these are present in the training data of RNA–X and four (AATF, FAM120A, HLTF and PHF6) are excluded from it entirely. For BAnG and RNAtranslator the latter four are not excluded, which gives them an advantage on exactly the targets where RNA–X operates zero-shot. We compare all designs against the known natural binders of each protein and against two negative controls: 100 randomly selected natural RNAs and a set of experimentally validated non-binders.

As shown in figure 2C, the score distributions for ELAVL1 are comparable for GenerRNA, RNAtranslator and RNA–X, all three approach the distribution of known ELAVL1 binders and exceed RNAGen and BAnG. For RBM5, which has three orders of magnitude less interaction data, RNA–X outperforms all other methods, indicating its ability to generalize in data-scarce settings. For the six seen targets in Figure 2A, RNA–X obtains a higher median predicted binding score than both BAnG and RNAtranslator on N of them. For the four excluded targets (Figure 2B), RNA–X matches or exceeds RNAtranslator and outperforms BAnG on all four, despite being the only method evaluated in a true zero-shot setting. The five remaining seen targets follow the same trend (Supplementary Figure 3B). To test whether RNA–X generalizes beyond the benchmarked set, we additionally designed RNAs for 71 further proteins and RNA–X surpasses the random-RNA control on 50 of these 71 targets (Supplementary Figures 11–12), indicating that the model transfers broadly across the RBP landscape.

**Fig. 2:**
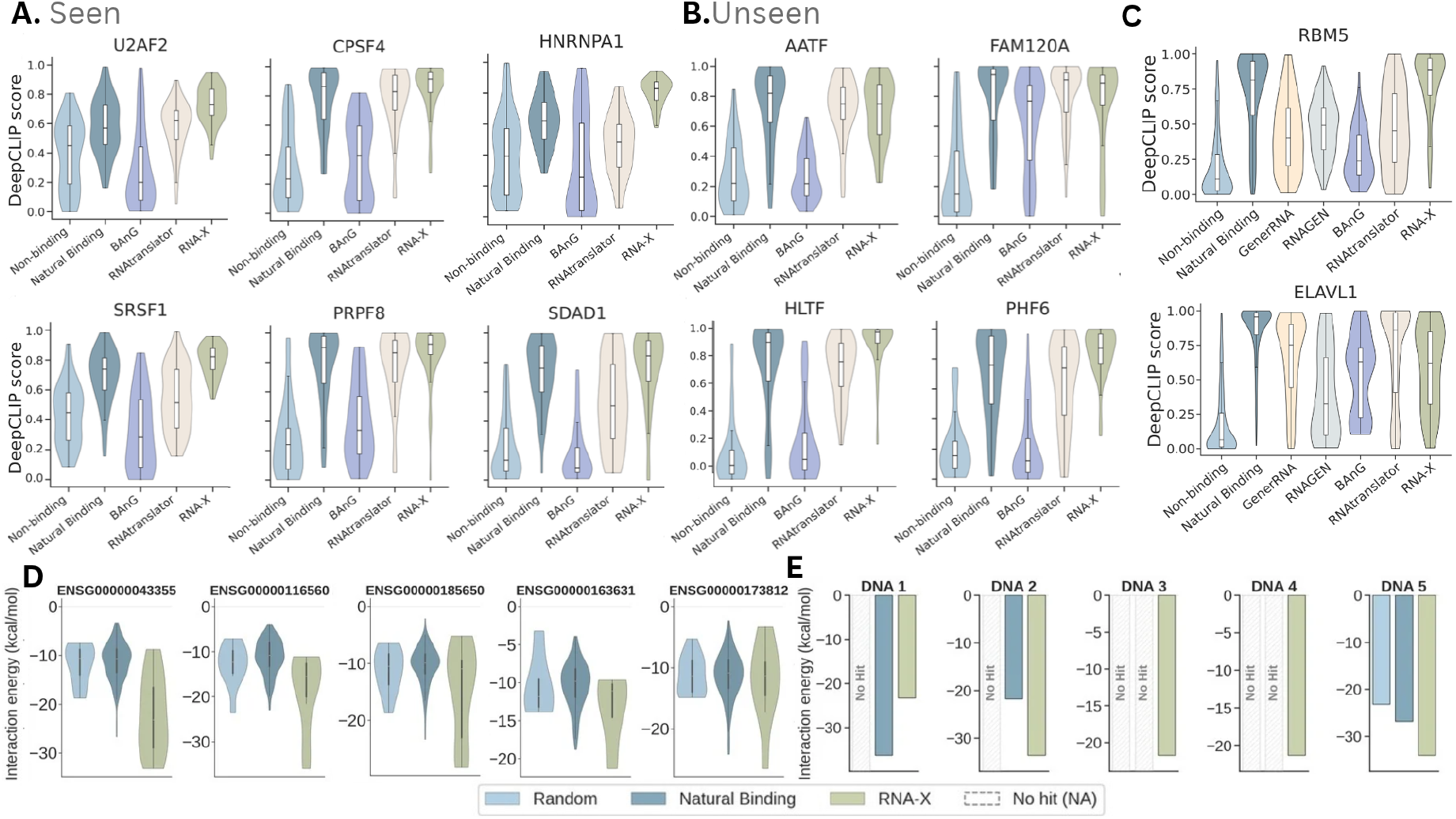
Comparison of RNA design methods on protein targets and Unseen targets. In (A)–(C), scores are produced by the target’s own DeepCLIP model, and designs are compared against known natural binders and two negative controls (random natural RNAs and validated non-binders). **(A)** Six protein targets present in the training data of RNA–X. **(B)** Four protein targets excluded entirely from the training set of RNA–X, but not from those of BAnG or RNAtranslator and RNA–X is the only method evaluated zero-shot on these targets. **(C)** RBM5 and ELAVL1 are the only two targets on which RNAGen and GenerRNA are benchmarked, as both require per-target fine-tuning. **(D)** Designed RNA show low binding energy with target RNAs **(E)** Designed RNAs bind DNA targets and cases where no stable binding was detected are shown as NA.

The scoring and exploration stage drives much of this performance (Supplementary Figure 4). In an ablation that removes it, generating with iterative unmasking alone (RNA–X *no exploration*), the mean predicted binding score drops across the benchmarked targets (Supplementary Figure 4A), the novelty rate falls from 72% to 59% (Supplementary Figure 4C), the full model reaches a lower minimum free energy per nucleotide than the natural binders themselves (Supplementary Figure 4D), and sequence diversity declines as designs collapse onto fewer distinct clusters (Supplementary Figure 4B). Exploration therefore improves binding, novelty, stability and diversity together, rather than trading one against another. Overall, RNA–X designs are novel with respect to the training data, fold at least as stably as natural binders, and remain sequence-diverse, with full analyses in Supplementary Note 1.10.

### 2.3 RNA–X can design binding RNA for unseen RNA and DNA targets

We next evaluate RNA–X on RNA and DNA targets whose interactions were not included during training. Since no competing method can design RNA against these target types, we compare our designs with known natural binder RNAs and with random natural RNAs as negative controls. For RNA targets we select those with the largest number of known binding RNAs, so that the comparison against natural binders is statistically meaningful. Each DNA target in the test set has only a single known binding RNA, so DNA targets are chosen at random. For every target we design as many RNAs as there are natural binders. Binding is predicted with IntaRNA [28] for RNA–RNA and RIsearch [47] for RNA–DNA interactions (Supplementary Figure 3A).

Figure 2D shows the results for five RNA targets. The designed RNAs show strong binding potential, with interaction energies comparable to or lower than those of the natural binders, indicating that the designs adopt structures favourable for interaction. Figure 2E shows five DNA targets. Here the predicted binding energies of our designs are more favourable than those of the natural binders for two targets and slightly worse for one. For the remaining two, RIsearch detected no stable interaction for either the random or the natural binder RNAs (shown as *No Hit*), whereas our designs achieved negative interaction energies, indicating stable binding where no natural reference could be scored.

We extend this evaluation to a wider set of unseen targets in Supplementary Figure 3C,D, covering three additional RNA and six additional DNA targets. The same trends hold: RNA–X designs reach interaction energies at or below the natural binders on the majority of these targets, and produce stable interactions for several DNA targets where the controls yield no hit.

### 2.4 RNA–X-designed RNAs silence an endogenous gene in human cells

To test whether RNA–X designs are functional in cells, we validated the RNA–RNA setting experimentally. We conditioned RNA–X on the 5′ untranslated region (5′UTR) of the human androgen receptor (*AR*) mRNA, the principal driver of prostate cancer progression and a long-standing therapeutic target [46]. The 5′UTR governs translation of the downstream coding sequence, so an RNA engaging it should reduce AR protein, and the design requires no protein-level information about the target.

#### Design and setup

We generated four RNA sequences (RNA1–RNA4) in a single design run, with no post-hoc filtering and no target-specific fine-tuning, and transfected them together with a random RNA control into 22Rv1 cells, which express AR endogenously. After 48 h, in three independent biological replicates, we measured *AR* mRNA and full-length AR protein (AR-FL), and also the transcript of *KLK3*, a canonical AR-regulated gene encoding prostate-specific antigen that reports AR transcriptional activity [9]. *KLK3* shares no complementarity with our designs, so a change there without a corresponding change in AR would signal a nonspecific response to transfected RNA. Sequences and full procedures are in Supplementary Note 1.13 and Supplementary Tables 5–8.

#### RNA–X designs reduce target mRNA and protein levels

RNA3 reduced *AR* mRNA to ~60% of the random control (*p <* 0.0005; Figure 3A) and AR-FL protein to ~57% (*p <* 0.005; Figure 3C,D). The other three designs produced reductions that did not reach significance (RNA1, RNA2) or no change (RNA4), and the control produced no effect. The mRNA and protein measurements agree in rank across all four designs, indicating that the effect of RNA3 is a property of that sequence rather than experimental variation.

**Fig. 3:**
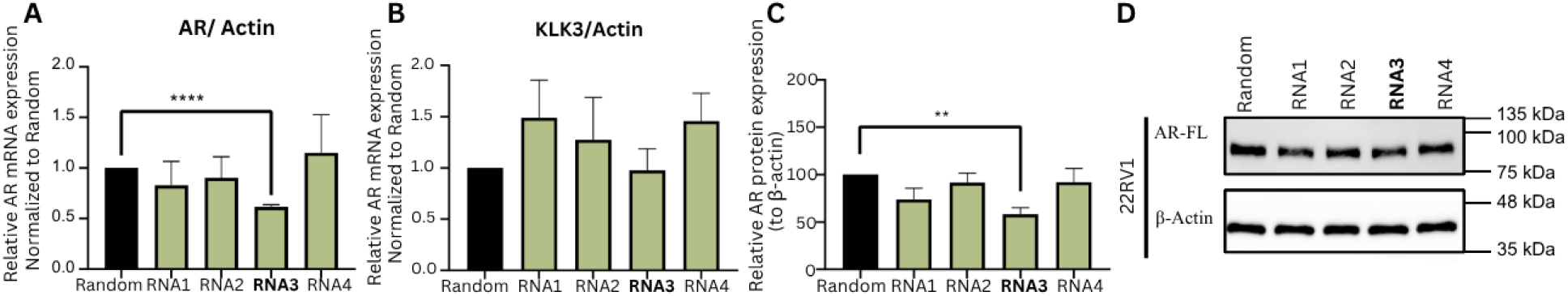
RNA–X-designed RNAs targeting the *AR* 5′UTR reduce AR levels in prostate cancer cells. RNA1–RNA4 were designed by RNA–X conditioned on the 5′UTR of the human *AR* mRNA; a random sequence served as the negative control. 22Rv1 cells were transfected with the in vitro transcribed RNAs and analyzed after 48 h. **(A)** *AR* mRNA levels quantified by qPCR, normalized to *β*-actin and shown relative to the random RNA group. **(B)** *KLK3* mRNA levels, a readout of AR transcriptional activity, quantified and normalized as in (A). **(C)** Quantification of AR-FL protein, normalized to *β*-actin and shown relative to the random RNA group. **(D)** Representative western blot of AR-FL and *β*-actin (loading control) in 22Rv1 cells transfected with the indicated RNAs. All bar plots show mean ± SEM of *n* = 3 independent experiments; multiple *t*-test against the random RNA group; **** *p <* 0.0005, ** *p <* 0.005, otherwise not significant.

#### Why RNA3?

Among the ten strongest IntaRNA-predicted interactions between the four designs and the *AR* 5′UTR, five belong to RNA3, against three for RNA1 and one each for RNA2 and RNA4 (Supplementary Figure 14). RNA3 engages five well-separated positions spanning the UTR, with energies between −18.50 and −16.37 kcal/mol and no weak site among them. The other designs depend more on a single dominant contact: RNA1 forms the strongest individual interaction (−21.50 kcal/mol) but with a wider spread across its remaining sites. The activity of RNA3 is therefore consistent with broad, uniform engagement of the target rather than a single high-affinity contact, though four designs are too few to establish this as a general predictor of cellular activity.

#### Downstream transcriptional output

*KLK3* was unchanged for all four designs (Figure 3B). That an unrelated transcript stays at control levels in samples where AR-FL is clearly reduced argues against a global effect of transfected RNA. The ~40% loss of AR-FL evidently falls below the threshold needed to deplete AR transcriptional output at this time point, which a dose–response series and later time points would be required to confirm. One of four sequences from a single, unguided generation round thus reduced both the transcript and the protein level of its target, with no screening library, enrichment cycle, or target-specific training data.

### 2.5 RNA–X designs RNAs which are predicted to bind to therapeutically important target molecules

We evaluate RNA–X on five therapeutically relevant targets: three structurally distinct proteins (p53, a nuclear transcription factor; thrombin, a secreted serine protease; and EGFR, a membrane-bound receptor tyrosine kinase), the HIV-1 5′UTR as an RNA target, and the HIV-1 3′ LTR as a DNA target. The two HIV-1 regions contain conserved structural domains essential for viral replication and are established targets for inhibitory aptamers [37,42]. The targets and the evaluation pipeline are described in Supplementary Note 1.9. We compare our designs with experimentally validated aptamers for each target [22,27,48,12,42,37] and, as negative controls, with randomly chosen natural RNAs or RNAs validated as non-binders [37].

Across all targets, the designed RNAs form stable stem–loop structures with paired stems and accessible loops at the predicted interaction sites, with normalized MFE values comparable to or lower than those of the validated aptamers. For p53, the designs show stronger MM/GBSA-predicted binding energies [34] and more hydrogen bonds after molecular dynamics simulation in OpenMM [10] than both validated aptamers and random natural RNAs (Figure 4A). Thrombin and EGFR follow the same trend (Supplementary Figure 5). For the HIV-1 RNA target, our designs adopt structures similar to the validated 16 nt aptamer and achieve comparable or stronger interaction energies under MM/GBSA and IntaRNA [28] (Figure 4B). For the HIV-1 3′ LTR, on both sense and antisense strands, the designs bind more strongly than the random baselines and perform comparably to validated DNA-targeting aptamers under RIsearch (Figure 4C), in which the random RNAs produce no detectable signal at all.

**Fig. 4:**
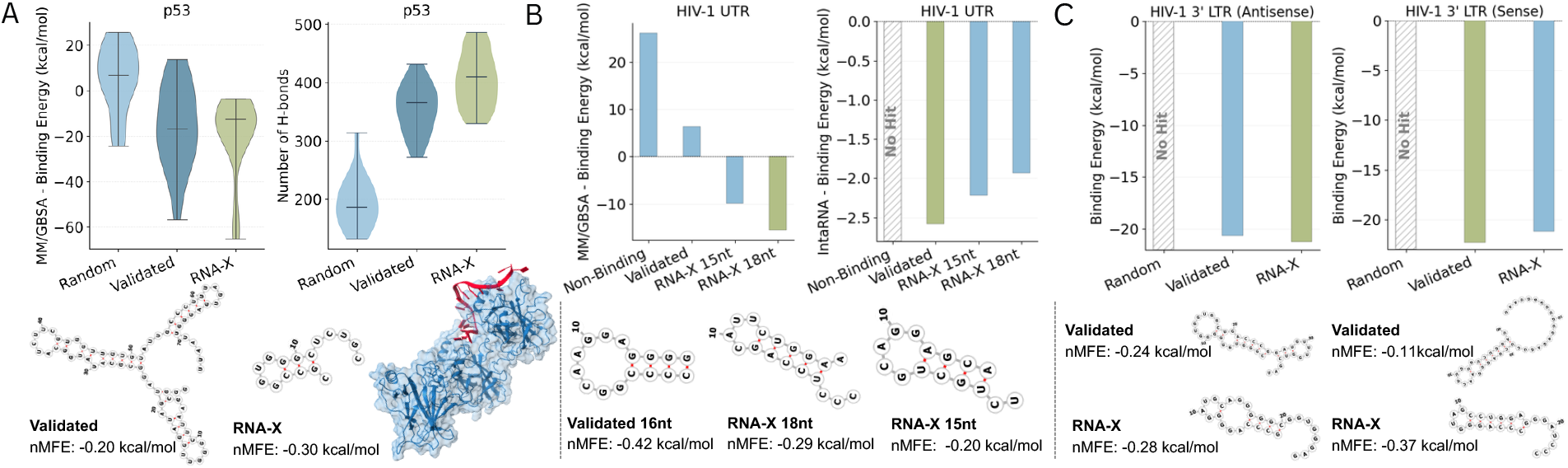
Evaluation of RNA–X on therapeutically relevant targets. Binding energy distributions, secondary structures of designs and controls, and the predicted 3D structure of the RNA–p53 complex. **(A)** p53: RNA–X designs show lower normalized MFE, stronger MM/GBSA-predicted binding energies and more hydrogen bonds during simulation than validated aptamers. **(B)** HIV-1 5′UTR RNA target: designs form stable stem–loop structures with MFE values comparable to or lower than validated aptamers. **(C)** HIV-1 3′ LTR DNA, sense and antisense strands: designs perform comparably to validated DNA-binding aptamers under RIsearch.

### 2.6 RNA–X learns meaningful representations across different interaction types and targets

To assess the representations learned by RNA–X, we analyze randomly sampled experimentally validated interactions from the test set at two levels. First, on a subset balanced across the three interaction types, a t-SNE visualization of RNA representations yields well-separated clusters by interaction type (Supplementary Figure 8A), showing that the model captures interaction-aware features. Second, within RNA–protein interactions, RNAs targeting proteins from the same cluster group together (Supplementary Figure 8B), indicating that the representations also encode fine-grained target-specific information.

We then evaluate the learned representations on three benchmark tasks (See Supplementary Note 1.14 for details). RNA–X achieves the highest AUROC and AUPRC on every dataset with low variance, matched only by AIDO.RNA on shRNA efficiency prediction (Supplementary Figure 7), despite having orders of magnitude fewer parameters (135M) than EVO2 (1B) and AIDO.RNA (1.6B). Removing the representation head lowers performance consistently (RNA–X *wo-head*), confirming that the geometry correction contributes beyond the pretrained backbone.

### 2.7 Multi-target RNA Design and Scaffold Optimization

We investigate whether RNA–X can one-shot design RNAs that bind multiple targets simultaneously. We use the Cas9–DNA–guide RNA complex (PDB 4OO8) [29], whose guide RNA divides into a DNA-complementary spacer and a Cas9-binding scaffold. We generate RNAs in three modes to show the model’s control over generation. The first is a complete RNA designed *de novo* from both the Cas9 protein and the target DNA (RNA-X-complete). The second is the scaffold alone designed *de novo* with the spacer held fixed (RNA-X-scaffold). The third optimizes the wild-type (WT) scaffold rather than rebuilding it, under a composite score that combines target binding, structural similarity to WT and thermodynamic stability (RNA-X-scaffold-opt). Full details are in Supplementary Note 1.8.

The three modes trade off differently against the WT (Supplementary Figure 9). RNA-X-complete gives the least favorable MM/GBSA binding energies at both interfaces. RNA-X-scaffold keeps strong RNA–DNA binding, as intended, but loses RNA–protein affinity. The scaffold-optimized designs recover the balance. They match or exceed the RNA–DNA binding energy of the WT while restoring RNA–protein binding, and all designs are more intrinsically stable than the WT by normalized MFE. Interface energetics, hydrogen bonds, secondary structures and a comparison of the WT and designed 3D complexes are reported in Supplementary Note 1.8 and Supplementary Figures 9 and 10. RNAGenesis [51] fixes the spacer and selects among a pool of scaffold point mutations. In contrast, RNA–X optimizes or designs novel RNAs without additional tools which enables choosing between conservative (RNA-X-scaffold), globally optimized (RNA-X-scaffold-opt) and fully novel (RNA-X-complete) designs.

The Cas9 system pairs targets of different types. To test multi-target design more systematically, we designed RNAs for all six pairwise combinations of the four proteins excluded from training (AATF, FAM120A, HLTF, PHF6). The dual-target designs score above both the random-RNA and the non-binding controls for every protein in every pair (Supplementary Figure 13). This shows that simultaneous conditioning is not restricted to the structurally coupled Cas9 case and extends to protein pairs the model has never seen.

## 3 Discussion

We introduce RNA–X, the first RNA interaction foundation model, which learns RNA interactions with proteins, RNA and DNA in a shared representation space. Unlike previous approaches restricted to a single interaction type, RNA–X designs RNAs for diverse molecular contexts and can be conditioned on multiple targets at once, indicating that it captures higher-order interaction principles rather than memorizing iso-lated patterns. RNA–X enables controllable RNA desing by adjusting scoring weights biases designs toward particular structural or biochemical constraints. Such control proves essential for Cas9 guide RNAs, where both multi-target binding and preservation of the native secondary structure are required. Representation learning matters equally. Without corrective losses, sequence embeddings collapse into narrow cones and enforcing alignment, whitening and uniformity restores isotropy and yields embeddings that transfer better to downstream tasks. Our cellular experiments show that these designs are functional, not merely plausible in silico.

Despite these strengths, several challenges remain. Although our computational assessments indicate strong binding and MD simulations provide dynamic insight into complex stability, these methods are inherently imperfect. Molecular dynamics can introduce errors due to initialization choices or boundary effects. Tools such as DeepCLIP offer valuable predictions, but they rely on protein–RNA datasets from specific RNA-binding proteins and therefore cannot be generalized to proteins without available training data. Moreover, our framework does not yet explicitly control for designing RNAs that bind at a precise site or motif within the target molecule.

## 4 Methods

### 4.1 Datasets and preprocessing

**Data**. We curate a dataset of ~102 million RNA interactions from databases listed in Supplementary Table 1. The data is filtered and clustered as described in Supplementary Note 1.1. Detailed statistics for interaction types and train/test splits are provided in Supplementary Tables 2, 3 and 4, respectively. An overview of the RNA and protein type distributions in the dataset is shown in Supplementary Figure 2. **Tokenization**. Each amino acid in target proteins and nucleotide in nucleic acids are represented by a single token, yielding a vocabulary of 35 tokens (See details in Supplementary Note 1.2). **Sampling**. Then, to reduce bias due to overrepresented molecules and hubs, we apply a static weighting scheme that penalizes frequent RNAs and targets while increasing the importance of underrepresented molecules. Each interaction is assigned a weight *w* = (*deg*_RNA_)^*a*^(*deg*_target_)^*b*^(*clust*_RNA_)^*c*^(*clust*_target_)^*d*^]^*−γ*^, where the parameters *a, b, c, d* and *γ* regulate the impact of the molecular interaction count (deg) and the size of the cluster (clust). Feature values are normalized by medians in the data set, cut in the 99.5*th* percentile, and adjusted adaptively based on their dispersion between interaction types. During training, we maintain balance between interaction types by random selection and dynamically controlling the ratio of computational to experimental data. A linear scheduler shifts the focus from computational data early in training to experimental data later for improved generalization shown in Supplementary Figure 1. Within each subset, samples are drawn with respect to their normalized weights using inverse cumulative sampling (See Supplementary Note 1.3).

**Preventing data leakage**. Our evaluation separates two distinct notions of leakage, and we control for both: (i) after clustering the dataset during preprocessing, we split it into training and test sets so that no target or its close variants appear in both. Then during evaluations, a protein is treated as *unseen* only if it has less than 30% sequence identity to every protein in the training set. Anything above that threshold is treated as *seen*. Since training a reliable evaluator for a protein consumes interactions that would otherwise be available for training RNA–X, only four proteins are held out as unseen. (ii) The models used to score our designs are trained and tested entirely independently of them. For each target protein we train a dedicated DeepCLIP model on that protein’s own interaction data, split 80*/*10*/*10 into training, validation and test sets, and report its AUROC on the held-out test set. The RNAs designed by RNA–X are never part of any DeepCLIP training set, and the natural binders we compare against are drawn only from the test split. Scores from these two sources are therefore directly comparable. We release all trained evaluator models, together with their data splits and test-set performance, so that every score reported in this work can be reproduced and so that the models can be reused as affinity predictors for the proteins they cover.

### 4.2 RNA–X Architecture

RNA–X is a transformer-based masked language model designed to jointly model RNA and its molecular targets. It consists of *L* pre-layer-normalized Transformer encoder layers, each composed of multi-head self-attention and SwiGLU feed-forward sub-blocks. Attention operates over the concatenated RNA–target sequence, enabling bidirectional information exchange across molecule types. Let *r* = (*r*_1_, *r*_2_, …, *x*_*n*_) be the RNA sequence and *t* = (*t*_1_, *t*_2_, …, *t*_*m*_) the target sequence. We concatenate the two to form a single input token sequence *s* = [*t*; *r*] of length *n* + *m*. Each token is represented by the sum of three embeddings—token, position, and residue type—forming the input 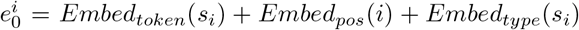 where 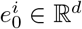 is the embedding of token *s*_*i*_. The embedding type of the residue distinguishes whether a token originates from RNA, protein, or DNA, allowing the model to capture biological context. At the final layer *L*, a linear projection and softmax produce nucleotide probabilities for each masked site, 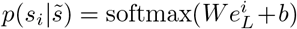 where W is the parameter matrix, b is the bias term and 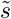 denotes a corrupted version of the input sequence obtained by replacing a subset of positions with a <monospace>[MASK]</monospace> token, the masking procedure is described in the following subsection. Training minimizes cross-entropy loss on masked tokens only, following the standard masked language modeling objective.

### 4.3 Pretraining by Unmasking

We model the problem as a conditional sequence modeling task, where both the RNA and the target sequence are presented as a single concatenated input to the model. To support both representation learning and conditional generation, we apply masking to both the RNA and the target tokens.

We apply random masking to the combined sequence *s* by selecting a subset of positions *M* ⊆ {1, …, *n* + *m*}. The model receives the corrupted sequence 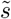, in which the tokens at positions *M* are replaced by a special <monospace>[MASK]</monospce>a token, and is trained to recover the original tokens at those positions. For all 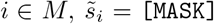, and for all 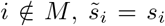. To jointly optimize sequence generation and later representation learning (see Section 4.4), we adopt a mixed masking strategy that combines structured and random corruption. For RNA sequences, in 80% of training steps the masking rate is drawn from a *β*(3, 9) distribution (mean ≈ 25%), and in the remaining 20%, it is uniformly sampled from [0, 1], resulting in an average mask rate of about 30%. This schedule allows the model to learn from both mild and severe sequence perturbations. For target sequences, a fixed 15% of residues are masked at random, providing a stable conditioning signal during training (See Supplementary Note 1.4 for masking details).

The objective is a single masked language modeling loss: 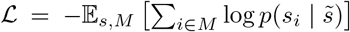. Since the model must predict tokens from both parts of the sequence using shared attention layers, it implicitly learns to model dependencies between RNA and target tokens. This results in embeddings that are sensitive to interaction-specific features rather than isolated sequence patterns.

### 4.4 Representation Learning

To learn stable and discriminative RNA–target embeddings, RNA–X uses a geometry-aware representation learning scheme. Although dynamic masking during pretraining produces contextual token embeddings, simple pooling often leads to modality-specific collapse, where RNA embeddings cluster with RNA and target embeddings cluster with target, creating anisotropic and low-rank structures. To avoid this, we apply a geometry-corrected projection head to the frozen encoder embeddings *e*^(target)^ and *e*^(rna)^, producing compact sequence vectors *z*_*t*_ and *z*_*r*_ via gated fusion and masked mean–max pooling (See the Supplementary Note 1.5 for details).

We train these representations with three complementary losses. (i) We apply a Barlow Twins objective. Let *z*_*t*_, *z*_*r*_ ∈ ℝ^*N* ×*d*^ where N is the number of samples and d is the representation size, to be the batch-standardized target and RNA embeddings, and let 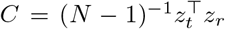 be the inter modality correlation matrix. The loss 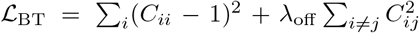 aligns positive RNA–target pairs while reducing feature redundancy; (ii) To prevent low-rank collapse, we apply the whitening regularization. For any standardized embedding *z*, we com pute *C* = (*N* − 1)^*−*1^*z*^*⊤*^*z* which is the intra-modality correlation matrix. We minimize 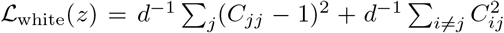 and apply it to both modalities: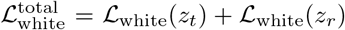 and (iii) To distribute embeddings evenly on the hypersphere, we use the uniformity loss 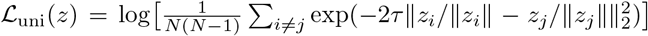 with *τ* = 2.0, applied to both modalities. The final objective combines these terms: 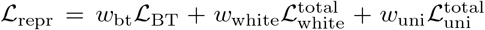, which jointly promotes alignment across modalities, isotropy within each modality, and uniform distribution across the representation space, resulting in balanced and interaction-aware embeddings.

### 4.5 Inference

At inference, RNA–X generates RNA sequences conditioned on a single or multiple target(s) through a three-stage process combining masked generation, refinement, and Monte Carlo Tree Search (See details in Supplementary Note 1.6). The model first performs iterative masked prediction, progressively filling masked regions of an RNA sequence. The generated RNA is then refined through mask–predict loops, where uncertain positions are re-masked and re-predicted until convergence. After each iteration, the candidate RNA is evaluated by a composite scoring function (See Supplementary Note 1.7 for details of scoring), and only the best-scoring RNA is retained for further refinement. This best-first procedure accelerates convergence and stabilizes optimization. Finally, Monte-Carlo Tree Search (MCTS) is applied for global optimization, where each node represents a candidate RNA sequence and edges correspond to single-base edits. Node selection follows the PUCT rule 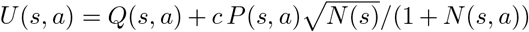, where *Q*(*s, a*) is the mean value of action *a* at state *s, P* (*s, a*) is the model-derived prior probability of choosing that action, *N* (*s*) is the total number of visits to node *s*, and *N* (*s, a*) is the number of times action *a* has been taken from *s*. The constant *c* controls the balance between the exploration of less-visited edits and the exploitation of high-value moves. Each expansion is guided by the nucleotide probabilities of the model and is immediately evaluated by the same scoring function, guiding the search toward RNA sequences with a predicted higher binding affinity, stability and structural quality.

