## Supplementary Material for "RNA–X: Modeling RNA interactions to design binder RNA and simultaneously target multiple molecules of different types"

### Supplementary Material for RNA–X: Modeling RNA interactions to design binder RNA and simulataneously target multiple molecules of different types

#### 1 Supplementary Notes

##### 1.1 Filtering and Clustering

To prepare our dataset, we first combined all available interaction datasets, since our model uses both the target molecule and the RNA sequence as input. In these datasets, most interactions include the full-length target and RNA sequences instead of only the binding sites. This leads to very long sequences and limited high-resolution information. To address this, we filtered the data so that the combined length of each target and RNA pair was less than 4096 residues. After filtering, we clustered the RNAs and targets using a 90% sequence identity threshold. For proteins, we used CD-HIT, and for RNAs, CD-HITest[21]. We performed clustering after merging all datasets of each interaction type. Clustering helped us remove duplicates and reduce oversampling of highly similar interactions. It also ensured that the model trained on diverse, representative samples and that closely related isoforms did not appear across training and validation sets. After filtering and clustering, we obtained comprehensive statistics for each interaction type, summarized in Supplementary Tables 2, 3, and 4. These tables show the diversity and size of our dataset, as well as the level of redundancy and overlap between experimental and computational sources.

##### 1.2 Tokenizing

To process sequence data, we design a custom residue-based tokenizer. Our tokenizer supports protein, RNA, and DNA sequences. We assign a unique token to each residue type. For example, we use "AA_A" for alanine in proteins, "RNA_G" for guanine in RNA, and "DNA_T" for thymine in DNA. In addition, we include special tokens [CLS], [EOS], and [MASK] to structure the input sequences and support masked language modeling. To represent unknown or uncommon residues, we use "AA_X", "RNA_X", and "DNA_X" tokens. In total, our tokenizer contains 35 tokens in its vocabulary.

##### 1.3 Sampling

To train our model effectively, we develop a sampling strategy that directly addresses the strong biases present in RNA interaction datasets. These biases occur because some targets and RNAs participate in many more interactions than others or they are highly overrepresented due to large cluster sizes. If we use uniform sampling, the model may overfit to these dominant patterns and neglect rare but important cases. To avoid this, we design a static weighting scheme for each interaction, with the goal of reducing the influence of frequent targets and large RNA clusters while promoting underrepresented examples.

Our weighting scheme combines several key features of each interaction: the degree of the RNA (i.e., the number of interactions it appears in), the degree of the target, and the cluster sizes of both RNA and target sequences. For each feature, we first normalize its value by dividing by the median of that feature across the dataset. To further limit the effect of extreme values, we clip all normalized features at the $99.5^{\text{th}}$ percentile. This normalization and clipping help to prevent numerical instabilities and ensure that no single sample can dominate the learning process.

The static weight for each interaction is then calculated as:

$$w = \frac{1}{\left[(\text{deg}_{\text{RNA}})^a \cdot (\text{deg}_{\text{target}})^b \cdot (\text{clust}_{\text{RNA}})^c \cdot (\text{clust}_{\text{target}})^d\right]^\gamma}$$

where $\text{deg}_{\text{RNA}}$ and $\text{deg}_{\text{target}}$ are the normalized and clipped degrees of the RNA and target, $\text{clust}_{\text{RNA}}$ and $\text{clust}_{\text{target}}$ are the normalized and clipped cluster sizes, $(a, b, c, d)$ are exponents that control the strength of each penalty, and $\gamma$ is a global smoothing factor.

To make the weighting adaptive, we automatically calculate the exponents $(a, b, c, d)$ for each interaction type and data source (computational or experimental) based on the observed dispersion (log-ratio of max to median) of each feature. Features with a larger spread receive higher exponents, meaning that outliers are down-weighted more aggressively. This approach ensures that the weighting is tailored to the specific imbalance present in each dataset (for example, RNA-protein targets can have up to 59,000 interactions, while RNA-DNA and RNA-RNA datasets are more balanced).

In addition to static weighting, we balance the sampling over interaction types and data sources. Our dataset consists of three interaction types—RNA-DNA, RNA-protein, and RNA-RNA—and includes both computational and experimental interactions. During sampling, we first select an interaction type randomly. Then, for each type, we use a linear scheduler to prioritize computational data early in training, gradually shifting focus to experimental data as training progresses. For each batch, we choose whether to sample from computational or experimental data according to the current weights from the scheduler. Finally, within each selected table, we sample individual interactions according to their normalized static weights using inverse CDF sampling. Supplementary Figure 1 illustrates the effect of our sampling strategy. As training progresses, we observe a balanced distribution of interaction types and a smooth transition from computational to experimental data. The application of static weights successfully reduces the dominance of large clusters and highly connected targets, resulting in a more equitable and representative training signal.

##### 1.4 Noise Schedule for Masking

To balance representation learning and sequence generation, we employ a mixed noise schedule during training. For RNA sequences, in 80% of training steps the masking rate is sampled from a Beta distribution $\beta(3, 9)$ (mean $\approx$ 25%), and in the remaining 20% it is sampled uniformly from $[0, 1]$. This combination yields an average overall mask rate of approximately 30% and exposes the model to a wide range of corruption levels, encouraging recovery from both mild and severe perturbations. For target sequences, a fixed 15% of residues are masked at random in every batch. This provides a consistent conditioning signal that helps stabilize the learning of target-conditioned representations. Together, these strategies balance structured and stochastic corruption, enabling the model to learn robust contextual embeddings and generate RNA sequences under varying levels of noise. This design is inspired by the dynamic masking schedule proposed in ESM-3 [13] and adapted here for multi-molecule RNA interaction modeling. A visual summary of the masking distributions is provided in Supplementary Figure 6.

##### 1.5 Representation Learning Head

We use the same training corpus, preprocessing, and sampling scheme as in pretraining. We freeze the pretrained backbone encoder used in pretraining. We train a geometry-corrected head that operates on frozen token embeddings. For each modality, we pass tokens through a small 1D-CNN stack with kernels, channels, GELU nonlinearity, batch normalization, and dropout. We fuse the backbone token embeddings with the CNN tokens using a gated residual adapter. Let $f_t^m \in \mathbb{R}^{T\times C}$ be frozen backbone tokens projected to the CNN width and $c_t^m \in \mathbb{R}^{T\times C}$ be CNN tokens for each modality $m$ (RNA or target). We compute a gated update $\Delta^m = \mathrm{FF}(\mathrm{LN}(f_t^m))$ and a gate $g^m = \sigma(W f_t^m)$. The fused tokens are

$$F_t^{\mathrm{m}} = \mathrm{LN}\big(c_t^m + \alpha\,(g^m \odot \Delta^m)\big)\,,$$

with a small learnable $\alpha$ (init 0.1). We instantiate one stream per modality (target or RNA). Then to obtain fixed-dimensional vectors for the RNA and the target, we apply masked mean-max pooling. Given a binary attention mask $a^m \in \{0,1\}^{T_m}$ indicating valid positions, we compute

$$\mu^m = \frac{\sum_i a_i^m F_i^m}{\sum_i a_i^m}, \qquad \eta_k^m = \max_{i:a_i^m=1} F_{i,k}^m,$$

and concatenate them:

$$u^m = [\,\mu^m;\,\eta^m\,] \in \mathbb{R}^{2C}.$$

Finally, modality-specific projection heads map these pooled features into the shared representation space:

$$z_t = f_{\mathrm{tar}}(u^{(\mathrm{target})}), \qquad z_r = f_{\mathrm{rna}}(u^{(\mathrm{rna})}).$$

These vectors, $z_t$ and $z_r$, are the target and RNA embeddings used in the representation learning objectives.

##### 1.6 Inference

At inference time, RNA–X generates RNA sequences conditioned on a given target molecule. Compared to one-directional autoregressive models, our iterative masked approach provides greater flexibility and control over RNA design and refinement. A design is produced in two stages: a *refinement* stage that performs iterative masked generation followed by mask–predict hill climbing, and a *search* stage that applies MCTS optimization[18] seeded by the stage-one design. Both stages use the same configuration, random seed and scoring function, so the only difference between them is the search itself; this is the ablation reported as RNA–X *no exploration* versus RNA–X. The target is conditioned as a sequence alone, so designing for a new target requires no interaction data, no known binders and no panel of related targets.

From the target sequence, and optionally a partial or template RNA, RNA–X initializes a masked RNA region of predefined length. The model repeatedly predicts masked tokens and fills them through iterative unmasking, guided at each step by the confidence (the maximum softmax probability) and entropy ($H(p) = -\sum_i p_i \log p_i$) of the model's token logits, optionally traded off with a small coefficient $\alpha$ (score = conf $-\alpha$·entropy). Sampling from logits is performed using configurable decoding strategies: greedy, multinomial sampling, top-$k$, or nucleus (top-$p$) sampling, with temperature-scaled probabilities. By default, we use top-$p = 0.75$ and temperature = 2.

The generated RNA is refined through multiple re-masking and re-prediction rounds that progressively improve binding and structural stability. In each refinement step, uncertain positions are re-masked based on low confidence or high entropy. The initial mask ratio is 0.75, which decreases by a rate of 0.001 per round until reaching a minimum of 0.01 or until the confidence threshold (0.9) or maximum score (0.95) is achieved. After each refill, we evaluate the candidate RNA with the scoring function described in Supplementary Note 1.7; if its score improves upon the current best, we accept it and continue refinement from this new best. This best-first update accelerates convergence and reduces unnecessary exploration, while randomized masking (`epsilon`= 0.2 or Gumbel with $\tau = 1.0$) avoids local optima. Masked regions are refilled using the same iterative generator but with reduced temperature (1.0) and narrower sampling thresholds. This loop continues for up to 10,000 refinement rounds or until convergence.

Raw head logits are not comparable across targets or design lengths, so the binding term of the scoring function is normalized against a reference pool built from the generator itself. Before the first design at a given length, a single forward pass on a fully masked sequence yields the generator's per-position marginals, from which 300 sequences are sampled independently and scored. We take $\mu$ as the median logit of this pool and $\tau$ as its interquartile range divided by 1.35, floored at 1.0, and normalize each candidate's logit as $(\text{logit} - \mu)/\tau$. Median and interquartile range are used rather than mean and standard deviation because the pool reaches very large logits. The pool's median base entropy, median GC content and median mean-field negative log-likelihood are stored as reference values for the corresponding penalty terms. Every reference value is therefore the generator's own rather than that of a natural binder, which is what allows the procedure to run on targets with no known interaction partners. The pool is cached per distinct design length.

To explore the sequence space beyond local refinements, we employ Monte-Carlo Tree Search (MCTS) guided by model logits and scoring feedback. Each node represents an RNA sequence, and edges correspond to single-base edits. The prior for each edit action is derived from the model's nucleotide probabilities, with the top-scoring positions ($K_{\text{pos}} = 28$) and bases ($K_{\text{base}} = 4$) retained. Identity edits are excluded to avoid redundant transitions. Node selection follows the PUCT rule:

$$U(s,a) = Q(s,a) + c \cdot P(s,a) \cdot \frac{\sqrt{N(s)}}{1 + N(s,a)},$$

where $Q(s,a)$ is the mean action value, $P(s,a)$ is the prior probability, $N(s)$ is the number of visits to state $s$, $N(s,a)$ is the number of visits to edge $a$, and $c$ $(= 1)$ controls the exploration–exploitation balance. Expansion and evaluation continue for 5,000 iterations or until a depth of 80 edits is reached. Each newly generated RNA is scored by the prediction model, and the best-valued sequence is retained as the final design.

##### 1.7 Scoring

For each generated RNA, binding probability is predicted by an interaction prediction module that inputs learned representations of the target and RNA sequences. Specifically, we train an interaction prediction head on top of the pretrained representation learning module of RNA–X to estimate RNA–target binding probabilities.

A separate head is trained for each interaction type—RNA–protein, RNA–RNA, and RNA–DNA—using the representations obtained from the representation learning head and frozen backbone. For each interaction type, we sample negative pairs by randomly pairing RNAs and targets that are not labeled as interacting in the dataset and train a binary classifier on the resulting positive and negative sets. Because this head uses the same training data employed during pretraining and representation learning, there is no data leakage between training and validation. Furthermore, since our evaluation relies on external predictors such as DeepCLIP[12], which are trained independently, there is no leakage between the model training and evaluation phases. The representation head is lightweight and trained only on the frozen representations, allowing it to capture binding-relevant relationships directly from the learned embedding space. Once trained, the head predicts a binding probability for any given RNA–target pair, serving as a key component of the sequence scoring process during generation.

Each generated RNA is then evaluated using a hybrid scoring function that integrates multiple criteria: (i) the predicted binding probability from the representation head and (ii) a set of structural, physicochemical, and heuristic penalties that account for RNA stability, composition, and potential biochemical artifacts. This combined scoring framework enables the model to balance binding affinity and structural feasibility, producing RNA designs that are both thermodynamically stable and highly compatible with their intended molecular targets.

RNA secondary structures are predicted using minimum free energy (MFE) folding computed by the ViennaRNA package [22]. The normalized stability score (STAB) is derived either from length-specific $z$-score calibration against empirical baselines or from a heuristic per-nucleotide energy band (favorable range $-0.4$ to $-0.2$ kcal·mol$^{-1}$·nt$^{-1}$)[27]. Structural shape features—loop exposure, global compactness, and GC content—are summarized into a composite measure (SHAPE) combining unpaired-loop fraction, stem compactness, and balanced GC ratio (0.35–0.65)[23].

Penalty terms are applied for low complexity (e.g., long homopolymer runs), self-association or reverse-complementary pairing, presence of G-quadruplex motifs, ribozyme-like catalytic cores, and translation-related elements such as Kozak sequences, Shine–Dalgarno motifs, or long open reading frames. An immunogenicity penalty accounts for GU-rich segments, long double-stranded regions, and poly-U tracts. A veto system enforces hard constraints—any RNA exhibiting strong ribozyme topology, extreme self-dimerization ($\geq 14$ contiguous matches), or long homopolymers ($\geq 9$) is immediately rejected[7][23].

The final composite score is computed as:

$$\text{Final} = B^{\star} \cdot \big(w_S \cdot \text{STAB} + (1 - w_S)\big) \cdot \big(w_H \cdot \text{SHAPE} + (1 - w_H)\big) \cdot \text{SPEC},$$

where $w_S = 0.4$ and $w_H = 0.4$ control the relative influence of stability and shape, and SPEC represents the specificity factor after aggregating penalties ($\text{SPEC} = 1 - P_{\text{total}}$, capped at 0.6). This formulation favors sequences that balance high binding affinity with structural robustness and developability constraints.

##### 1.8 Designing RNA to bind multiple targets

To evaluate whether RNA–X can generate RNAs that interact with multiple molecular targets simultaneously, we test it on the Cas9–DNA–RNA ternary complex (PDB ID: 4OO8)[24]. This complex involves a guide RNA bound to both the Cas9 protein and its complementary DNA target. The guide RNA comprises two functional regions. The spacer sequence base-pairs with the DNA, and the scaffold region binds Cas9 and maintains the structural fold necessary for activity.

We first input the model with the Cas9 protein and its DNA target as conditioning sequences and initialized the RNA input as fully masked. The model then generates an RNA candidate intended to bind both targets simultaneously. We next analyze whether RNA–X could refine the Cas9-binding scaffold while preserving the DNA-targeting spacer. We fix the spacer region and mask only the scaffold region, which prompts the model to redesign the scaffold sequence while maintaining the spacer unchanged. This step tests the model's ability to contextually modify structural domains under mixed constraints.

Since the efficiency of guide RNAs (sgRNAs) depends strongly on their secondary structure, we change the scoring so that it preserves the native folding of the wildtype RNA while improving stability and binding toward multiple targets. The final optimization objective combines three components. These are target binding, secondary-structure similarity, and thermodynamic stability. For a candidate RNA sequence $r$, we define the total score as:

$$\text{Score}(r) = B(r) + S(r, r_{\text{wt}}) + \mathit{\Delta}\text{MFE}(r, r_{\text{wt}}),$$

where $B(r)$ is the average predicted binding probability of the RNA to all targets (Cas9 and DNA) obtained from the trained representation heads, $S(r, r_{\mathrm{wt}})$ measures the structural similarity between the candidate and wildtype RNA, and $\Delta\mathrm{MFE}(r, r_{\mathrm{wt}})$ quantifies the improvement in minimum free energy (MFE) stability. The last term is computed as the normalized energy difference between the candidate and wildtype RNA folds. This composite scoring ensures that redesigned RNAs preserve the structural features essential for Cas9 recognition, since sgRNA efficiency depends critically on its secondary structure, while still allowing improvements in binding and overall stability. By integrating multi-target binding scores with structural similarity and thermodynamic energy terms, the optimization balances biological fidelity with functional enhancement.

For each RNA (WT, spacer-fix, structure-opt-1/2/3, and complete) we predict the ternary Cas9–DNA–RNA complex using Boltz2[25]. The inputs are the Cas9 protein, the target DNA duplex, and the candidate guide RNA. We estimate binding energetics with UniGBSA. Specifically, we compute MM/GBSA interaction energies (kcal/mol) for the RNA–protein and RNA–DNA interfaces from the selected ternary structure. More negative values indicate stronger binding. The same protocol is applied to all variants to ensure fair comparison. For the pairwise protein experiments, a single RNA is designed per pair, conditioned on both protein sequences, and then scored independently with each protein's DeepCLIP model.

To quantify specific contacts, we count hydrogen bonds using MDAnalysis. We evaluate RNA–protein and RNA–DNA interfaces separately, using a donor–acceptor distance cutoff of 3.5 Å. Reported counts are per complex and directly comparable across designs.

We first compare the MM/GBSA interaction energies of all designs at both interfaces (Supplementary Figure 9A,B). *RNA-X-complete* gives the least favorable energies at the RNA–protein and the RNA–DNA interface. It also forms the largest number of hydrogen bonds with the protein (Supplementary Figure 9C), but these contacts do not translate into favorable energies. This suggests suboptimal contact geometry. *RNA-X-scaffold* keeps strong RNA–DNA binding, as intended by the fixed spacer, but loses RNA–protein affinity. The scaffold-optimized designs (*RNA-X-scaffold-opt* 1/2/3) recover the balance between the two interfaces. They match or exceed the RNA–DNA binding energy of the WT while restoring RNA–protein binding, and *RNA-X-scaffold-opt* 3 is the strongest on RNA–DNA.

Second, we compare the thermodynamic stability of the designs. All designs have a more negative normalized MFE than the WT (Supplementary Figure 9E). This indicates higher intrinsic stability.

Third, we inspect the predicted secondary structures (Supplementary Figure 9D). *RNA-X-scaffold* preserves the canonical spacer and makes only local edits to the scaffold. The scaffold-optimized designs form compact stem loops that strengthen the Cas9-interacting hairpins without losing spacer complementarity.

We then compare the 3D structures of the wild-type (WT) Cas9–DNA–RNA complex with all designed RNA sequences to understand how each design changes the protein–RNA interface. Protein residues that interact with RNA are identified using a 3.0 Å heavy-atom distance cutoff.

First, we measure the number of protein residues that contact RNA (Supplementary Figure 10A). The WT shows a moderate number of contacts. The *RNA-X-complete* design has the highest number of contacting residues, meaning that it forms many new interactions. The *RNA-X-scaffold* design has slightly fewer or similar contacts compared to the WT. The scaffold-optimized versions (*RNA-X-scaffold-opt* 1/2/3) recover many contacts and sometimes show even more than the WT.

Second, we compare how similar each designed binding site is to the WT using the Jaccard index, which measures how many contact residues are shared between two structures (Supplementary Figure 10B). The scaffold-optimized designs show the highest similarity to the WT binding site. The *RNA-X-scaffold* and *RNA-X-complete* designs share fewer WT contacts, meaning their interfaces shift away from the native one.

Finally, we look at which contacts are shared with the WT and which are new (Supplementary Figure 10C). The scaffold-optimized designs keep most of the WT contacts and introduce only a few new ones. The *RNA-X-scaffold* design keeps fewer WT contacts and adds several new contacts. The *RNA-X-complete* design introduces many new protein contacts, which explains its low similarity to the WT interface.

##### 1.9 Designing RNA to bind therapeutic targets

For protein targets we obtain the crystal structure from the Protein Data Bank [1] (1TUP [5] for p53, 4DII [26] for thrombin, 1NQL [11] for EGFR), removing any native ligands or aptamers. For the RNA target we take the sequence from the reference study and predict its structure with RNAComposer [2], since RhoFold is unreliable at this length. For DNA targets, AlphaFold3 complex predictions were low-confidence, so we report sequence-based energies only and compute no 3D structures or MM/GBSA values. We predict the structure of each design with RhoFold+ [28], model the complex with HDOCKlite [30] for proteins and HNADOCK [14] for RNA, and run molecular dynamics in OpenMM [10] (Supplementary Note 1.11). Binding energies are computed with UniGBSA [31], and hybridization energies with IntaRNA and RIsearch to confirm that predicted binding regions match those reported in the literature.

As shown in Supplementary Figure 5A, for EGFR, the designed RNAs achieve lower binding free energies and a higher number of hydrogen bonds compared to both validated and random RNAs, indicating stronger predicted binding and enhanced complex stability. Similarly, for Thrombin (Supplementary Figure 5B), our designed RNAs form well-structured stem–loop motifs with comparable or lower normalized minimum free energy (nMFE) values relative to validated aptamers, suggesting favorable folding and binding characteristics. Together, these results show that RNA–X is capable of generating RNA molecules with high predicted affinity and thermodynamic stability for therapeutic protein targets, demonstrating its potential for RNA-based drug design.

We designed RNA sequences to target two segments of the HIV-1 LTR: (i) the first 50 nt of the LTR for antisense-strand binding (comparable to B7) and (ii) the last region of the LTR for sense-strand binding (comparable to BS8). Validated aptamers from the two groups were included as positive controls. Because previous studies highlighted structural inaccuracies when predicting RNA–DNA complex structures computationally, and our AlphaFold3-based complex predictions showed low confidence, we did not compute 3D structures or MM/GBSA energies for these targets.

Instead, we analyzed RNA–DNA hybridization energies using the RIsearch tool, which is optimized for RNA–DNA pairing. We compared the predicted energies of our model's RNA designs with those of validated aptamers (B7 and BS8) and with random natural RNAs. The designed RNAs consistently exhibited binding strengths comparable to or better than the validated aptamers and were always stronger than random controls.

##### 1.10 Sequence novelty, stability and diversity

We characterize the designed RNAs along three axes. All measures are computed on 50 sequences per protein, drawn with a fixed random seed, and compared against the natural binders of each target, validated non-binders, random RNAs, and the ablated variant generated by iterative unmasking alone (RNA–X *no exploration*).

**Novelty.** To test whether RNA–X produces genuinely new RNAs rather than memorized variants, we compare each design against the all RNAs used for RNA–protein interaction training. Sequence similarity is screened with BLASTn [3]; to balance sensitivity across lengths we split candidates at 50 nt and set the word size to 4 for short and 7 for long sequences. A candidate is labelled *similar to known* if its best hit satisfies all of (i) $\geq 70\%$ identity, (ii) alignment length $\geq 15$ nt, (iii) query coverage $\geq 50\%$, and (iv) E-value $\leq 0.1$; otherwise it is counted as novel.

**Thermodynamic stability.** Each sequence is folded with RNAfold (ViennaRNA [22]) and we report the minimum free energy normalized by sequence length (nMFE), so that designs of different lengths can be compared directly. More negative values indicate more stable folds.

**Sequence diversity.** Each group is clustered with CD-HIT-EST [21] at eleven identity thresholds from 0.80 to 1.00, and we report the number of clusters per sequence. A value near 1.0 means that nearly every sequence forms its own cluster, while lower values indicate that sequences fall into shared clusters and are therefore more similar to one another.

Results for all five groups are shown in Supplementary Figure 4B–D.

##### 1.11 Molecular Dynamics Simulations

We performed molecular dynamics (MD) simulations using the OpenMM framework [9] to study the dynamic behavior of the designed RNA systems. All simulations were carried out under an isothermal–isobaric (NPT) ensemble at 310 K and 1.0 atm using a Langevin integrator. Each system was first energy-minimized and then equilibrated before entering a production run on CUDA-accelerated hardware. Systems with initial potential energies exceeding $10^6$ kJ/mol were subjected to extended minimization protocols of up to $10,000$ steps to ensure stability. All systems were solvated in a TIP3P water box with 1.0 nm padding and neutralized to 0.15 M ionic strength using $Na^+$ and $Cl^-$ ions. Prior to simulation, structures were preprocessed with PDBFixer [10] to correct missing atoms and residues and prepared using the AMBER14 force field with protonation states adjusted for pH 7.0. Trajectories were recorded at regular intervals for subsequent analysis.

##### 1.12 Experimental Setup

During training, we use a tokenizer that representing each residue or base as a single token with additional special symbols. The model is a Transformer encoder with $L = 14$ layers, hidden dimension $d_{\text{model}} = 768$, $n_{\text{heads}} = 12$, and maximum context length 4096. To improve efficiency, we use FlashAttention-2 [8], which provides exact attention computation with reduced memory overhead and better parallelization. Multi-head attention is computed jointly over the concatenated RNA–target sequence to enable full cross-domain context exchange.

Training uses the AdamW optimizer with learning rate $5 \times 10^{-5}$, warmup of 4,800 steps, and a total of 50,000 optimization steps. Each device processes a batch of 16 samples, with gradient accumulation of 32, giving an effective batch size of $B_{\text{eff}} = B_{\text{device}} \times \text{world\_size} \times \text{grad\_accum}$. Training is conducted on two NVIDIA L40S GPUs in distributed mode. End-to-end training takes approximately one month under this setup. At inference time, generating a 28-nucleotide RNA sequence using the iterative masked decoding and refinement process requires about one minute on the same hardware.

##### 1.13 Wet lab Experimental Validation in Human Cells

**Target selection and design settings.** We used the $5'$ untranslated region ($5'$UTR) of the human androgen receptor (*AR*) mRNA, retrieved from Ensembl (transcript AR-201, ∼1,200 nt), as the conditioning target. The target was tokenized as an RNA sequence and the RNA input was initialized as fully masked with a fixed design length of 50 nt. Generation followed the standard three-stage procedure described in the Inference note and Supplementary Note 1.7, using the RNA–RNA interaction head for the binding term of the composite score, with default decoding settings (top-$p = 0.75$, temperature $= 2$ for initial unmasking; temperature $= 1.0$ during refinement). Four candidates were taken from a single design run. We applied no post-hoc filtering with an external affinity predictor, no re-ranking by a third-party tool, and no target-specific fine-tuning; the four sequences carried forward to the wet lab are the four top-scoring outputs of that run.

**Sequence and binding properties.** All four designs are 50 nt long with GC contents between 60% and 64%. Minimum free energies, computed with RNAfold (ViennaRNA [22]) at 37°C, range from −18.50 to −11.50 kcal/mol (−0.37 to −0.23 kcal/mol/nt). Hybridization energies with the target $5'$UTR were computed with RNAup [22], which accounts for the accessibility of both interaction partners. All four designs bind substantially more strongly than the random control (−21.50 to −18.07 kcal/mol versus −8.60 kcal/mol), and each forms five predicted binding sites on the target. Notably, the four designs converge independently on two regions of the 1,200 nt UTR, around positions 300–450 and 600–670, indicating that the model locates a small number of accessible sites rather than arbitrary complementary stretches. Sequences, oligonucleotides and all derived properties are listed in Supplementary Table 5.

**Negative control.** The control is a random RNA sequence of 40 nt (GC 40%, MFE −2.10 kcal/mol), with a predicted hybridization energy to the *AR* $5'$UTR of −8.60 kcal/mol. We note that the control is shorter and substantially less structured than the designs, so it controls for sequence composition rather than for length or fold.

**Off-target assessment.** To assess specificity, each design was screened against human RefSeq transcripts with BLASTn [3] and hybridization energies were computed for each hit. The number of predicted off-target transcripts varies considerably across the four designs: RNA1 (12 transcripts, 13 sites), RNA2 (20, 23), RNA3 (6, 6) and RNA4 (2, 2). For some designs, individual off-target hits reach hybridization energies more favourable than the on-target interaction. This screen is deliberately permissive: it considers sequence complementarity over the entire transcriptome without accounting for transcript expression in the cell line used, subcellular localization, or target accessibility, all of which constrain which interactions can occur in practice. It therefore bounds the possible off-target space rather than predicting it, and the cell-based measurements below are the relevant test of functional specificity.

**In vitro transcription.** Templates were ordered as complementary DNA oligonucleotide pairs (Macrogen Europe, order EO00006139, 0.05 $\mu$mol scale, MOPC purification) carrying a T7 promoter (`TAATACGACTCACTATA`) at the 5′ end of the forward strand (Supplementary Table 6). Forward and reverse oligos were incubated at 95°C for 5 min in annealing buffer (100 mM potassium acetate, 30 mM HEPES-KOH pH 7.4, 2 mM magnesium acetate) and cooled to room temperature. In vitro transcription was performed with the NEB HiScribe T7 High Yield RNA Synthesis kit according to the manufacturer's instructions, and the transcribed RNAs were purified as described below.

##### 1.14 Representation analysis

We evaluate the learned representations on three benchmark tasks spanning two interaction types: CRISPR on-target prediction (RNA–DNA) and siRNA and shRNA efficacy prediction (RNA–RNA). CRI-On [6] contains ∼15,000 sgRNAs targeting 1,071 genes across four human cell lines, with editing efficiency as the target. The siRNA datasets are Takayuki (702 sequences) [29] and Huesken (2,361 sequences) [15], both with binary labels. The shRNA dataset (2,076 sequences) provides efficiency scores [32]. We report AUROC and AUPRC for classification and Pearson and Spearman correlations for regression. For EVO2 we use the 1-billion-parameter pretrained version, the closest in size to our setup that remains practical to run. All foundation models are evaluated with frozen backbones and identical MLP classifiers trained with the same hyperparameters.

#### 2 Supplementary Figures

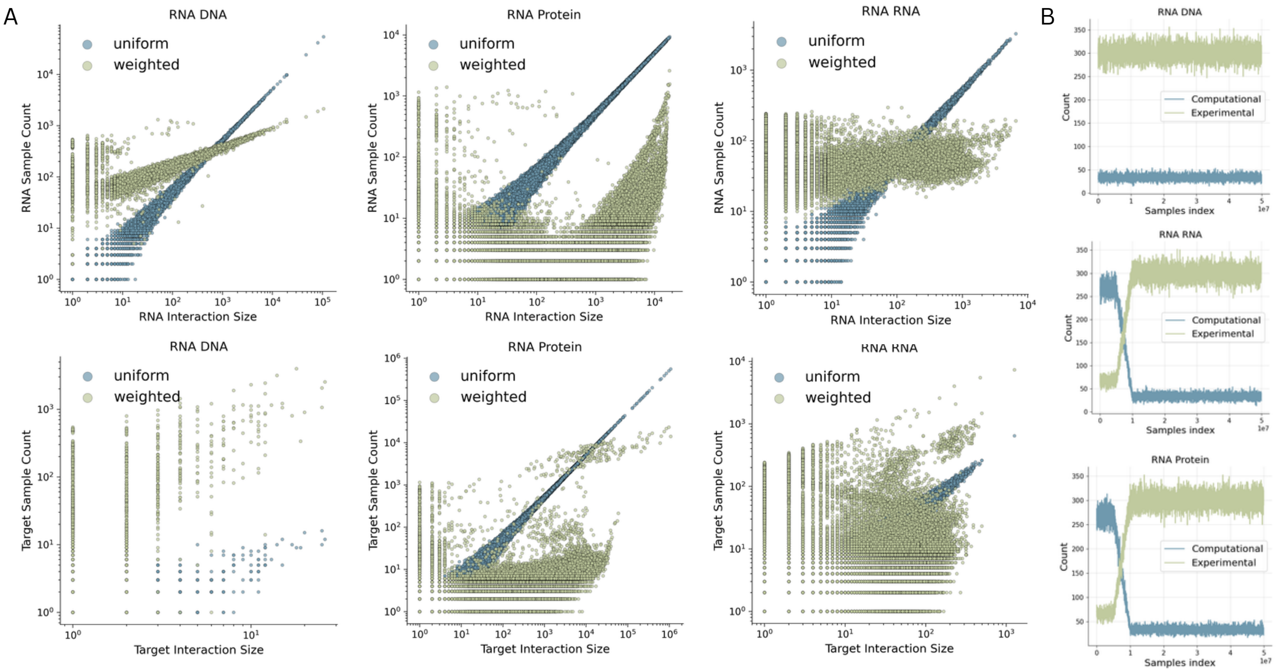


Supplementary Figure 1: **Sampling strategy for balanced model training. (A)** Comparison between uniform and weighted sampling distributions across RNA–DNA, RNA–protein, and RNA–RNA interaction datasets. Each scatter plot shows the relationship between the number of interactions per RNA (top) or per target (bottom) and their corresponding interaction sizes. Uniform sampling overrepresents frequent targets and RNAs, whereas our weighted sampling scheme significantly reduces this imbalance by down-weighting highly connected nodes and up-weighting rare interactions. **(B)** Sampling dynamics during training. The scheduler ensures balanced sampling across interaction types and gradually transitions from computationally derived interactions to experimentally verified ones.

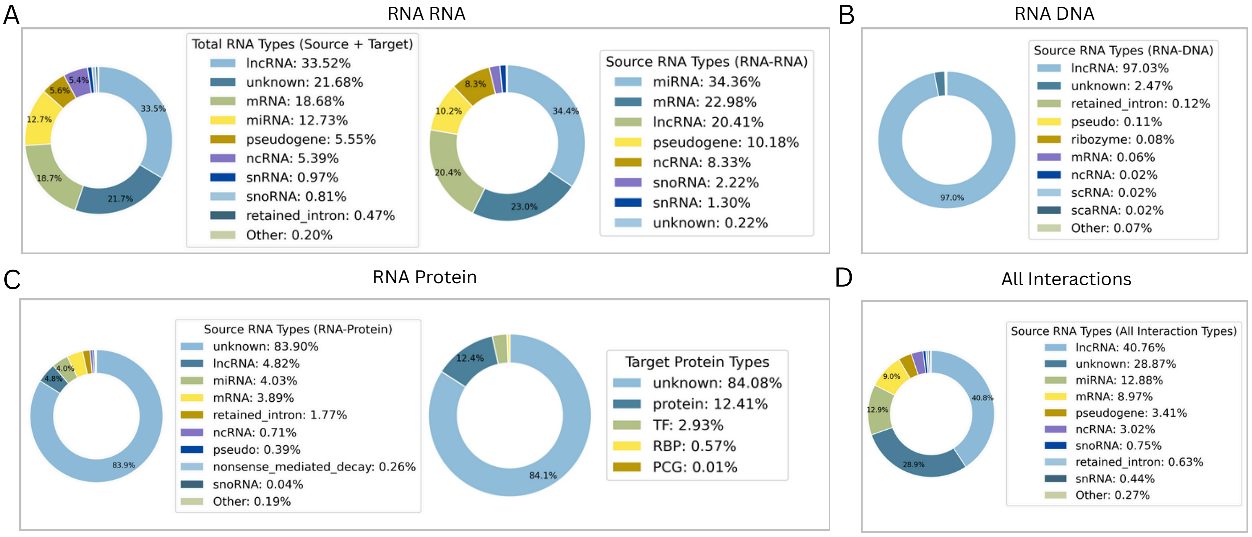


Supplementary Figure 2: **Analysis of RNA and protein types in our dataset** We include type information whenever it is available. The label unknown refers to RNAs or proteins that do not have type annotations in the original datasets. **(A)** RNA types in RNA–RNA interactions, shown for both all RNAs (source + target) and for source RNAs only. **(B)** RNA types used as source RNAs in RNA–DNA interactions. **(C)** RNA and protein types in RNA–protein interactions, including source RNA types and target protein types. **(D)** RNA types used as source RNAs across all interaction types in the dataset.

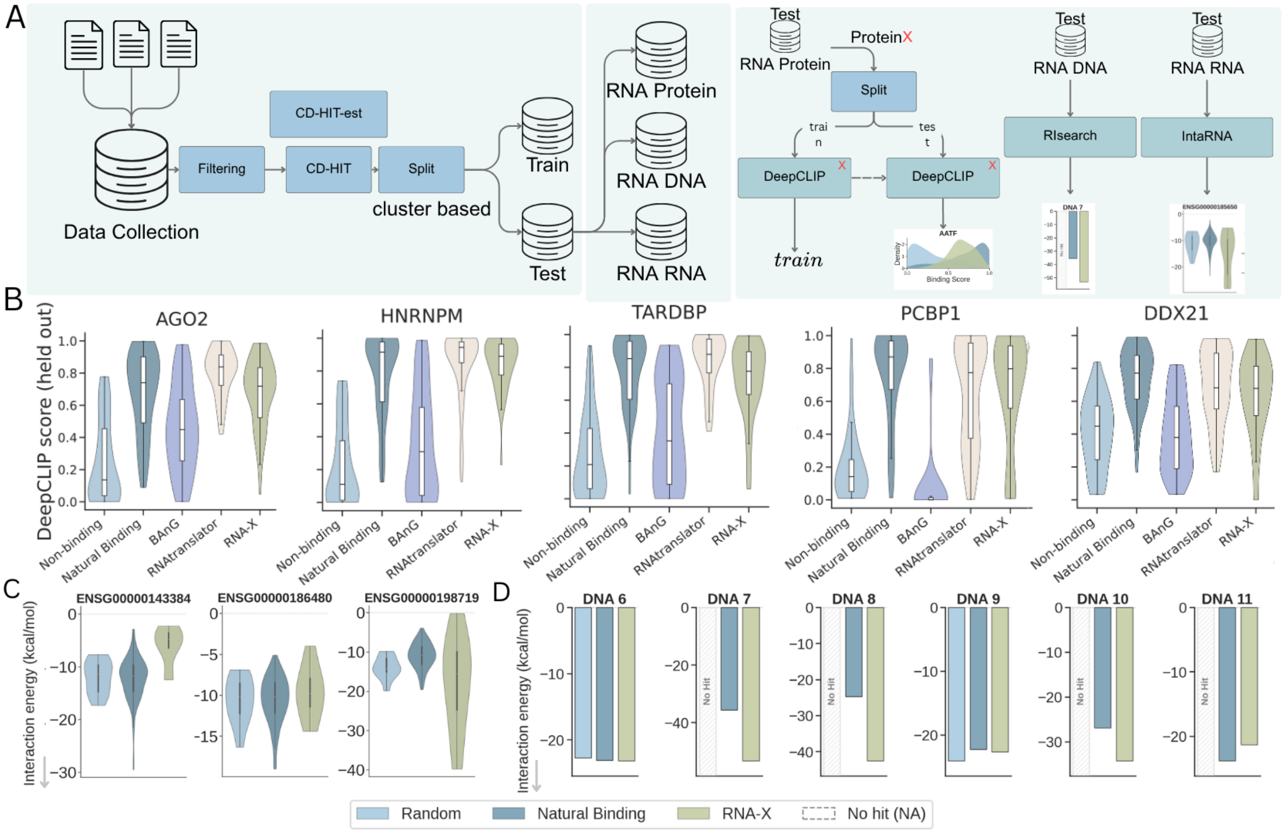


Supplementary Figure 3: **Data processing pipeline and additional target evaluations. (A)** Dataset construction and evaluation setup. Interaction data from all source databases are merged, filtered, and clustered with CD-HIT and CD-HIT-EST, then split cluster-wise into training and test sets and separated by interaction type (RNA–protein, RNA–DNA, RNA–RNA). For a held-out protein $X$, the RNA–protein test data are split again to train and evaluate a dedicated DeepCLIP$_X$ model, which is used only to score designs and never sees them during training. RNA–DNA and RNA–RNA designs are scored with RIsearch and IntaRNA, respectively. **(B)** The five remaining protein targets present in the training data of RNA–X (AGO2, HNRNPM, TARDBP, PCBP1, DDX21), complementing the six shown in Figure 2A. Designs are compared against known natural binders, validated non-binders, and the designs of BAnG and RNAtranslator, and are scored with each target's own DeepCLIP model. **(C)** Three additional unseen RNA targets, with interaction energies predicted by IntaRNA. **(D)** Six additional unseen DNA targets, with interaction energies predicted by RIsearch. In (C) and (D), lower (more negative) values indicate stronger predicted binding; bars marked *No Hit* indicate that no stable interaction was detected for that sequence group.

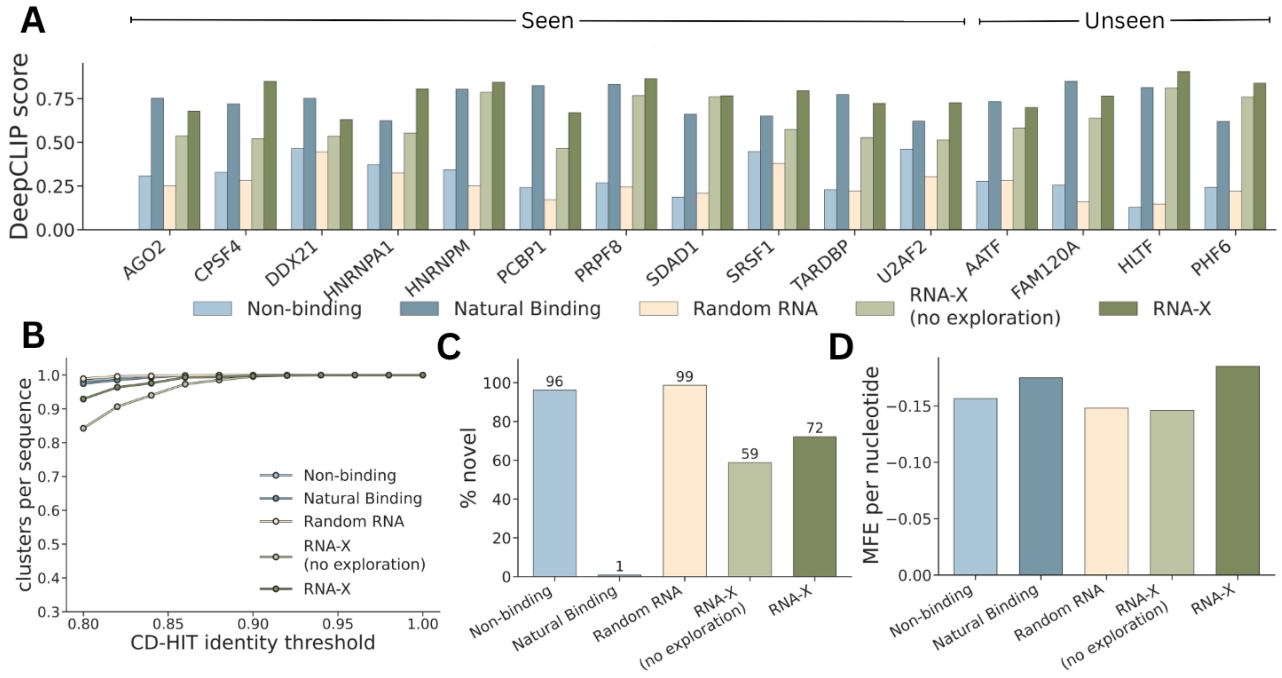


Supplementary Figure 4: **Per-target performance and characterization of RNA–X designs across the fifteen benchmarked protein targets. (A)** Mean predicted binding score for each target, ordered as the eleven proteins present in RNA–X's training data (Seen) and the four excluded from it (Unseen). Designs are compared against known natural binders, validated non-binders, randomly selected natural RNAs, and an ablated variant generated by iterative unmasking alone (RNA–X *no exploration*). **(B)** Sequence diversity, measured as CD-HIT clusters per sequence across identity thresholds; values near 1.0 indicate that nearly every sequence forms its own cluster. **(C)** Novelty rate with respect to the training corpus. **(D)** Minimum free energy per nucleotide (more negative indicates a more stable fold). In (B)–(D), values are pooled over all fifteen targets.

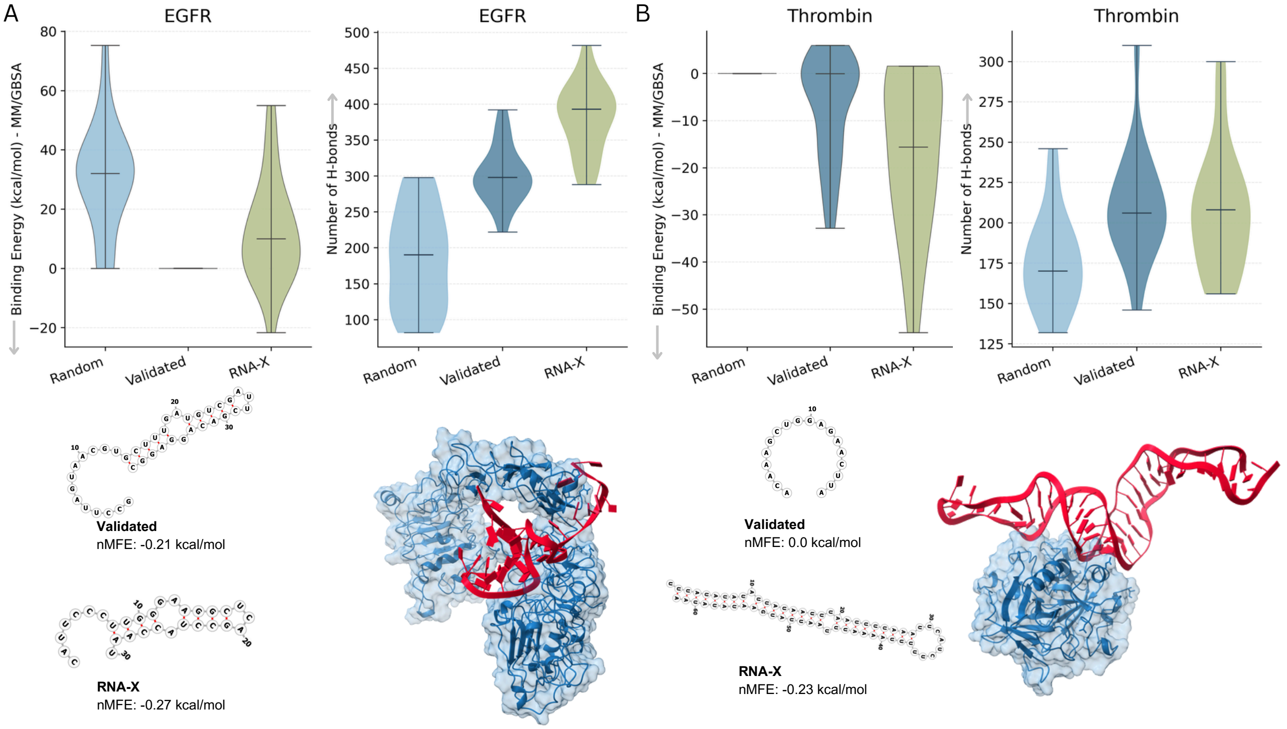


Supplementary Figure 5: **Evaluation of RNA–X-generated therapeutic RNAs targeting EGFR and Thrombin. (A)** Results for the EGFR protein target. Violin plots show the distributions of MM/GBSA-predicted binding free energies (left) and the number of hydrogen bonds formed after molecular dynamics simulations (right) for random RNAs, validated aptamers, and RNA–X-designed RNAs. Predicted RNA secondary structures are shown below each plot, along with their normalized minimum free energy (nMFE) values. The modeled RNA–protein complexes from molecular dynamics simulations (performed using OpenMM) demonstrate that RNA–X-generated RNAs adopt stable stem–loop conformations with multiple hydrogen bonds at the interaction interface. **(B)** Results for the Thrombin protein target, following the same evaluation procedure. Designed RNAs show lower or comparable nMFE values and form stable interactions with Thrombin, maintaining multiple hydrogen bonds within the binding interface. Across both therapeutic targets, RNA–X-generated RNAs exhibit stronger or comparable binding energies, well-structured RNA folds, and stable predicted complexes relative to validated aptamers, confirming their potential for therapeutic RNA design.

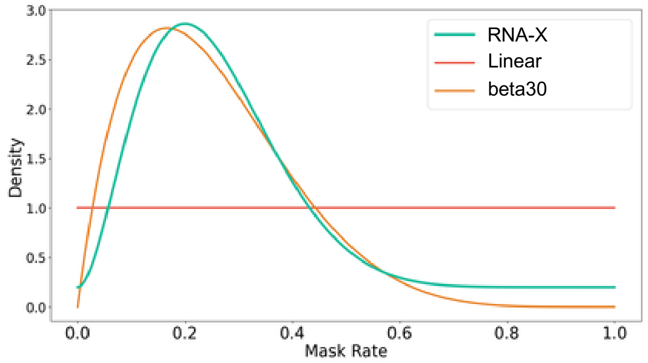


Supplementary Figure 6: Masking rate distribution used during training. A mix of Beta(3, 9) and uniform noise allows the model to handle various levels of corruption.

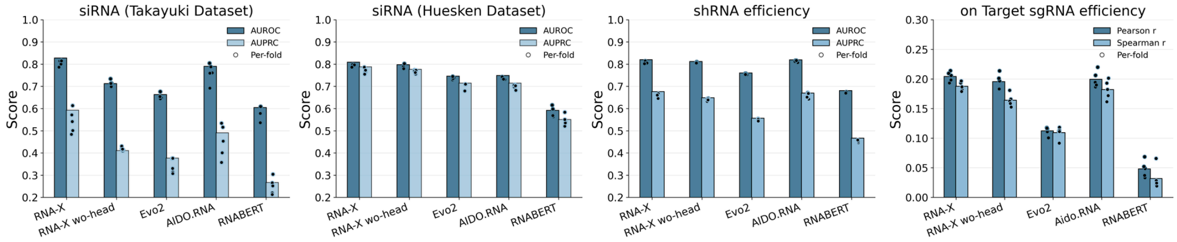


Supplementary Figure 7: **Representation learning and downstream performance of RNA–X.** Benchmark results on siRNA, sgRNA and shRNA efficacy prediction tasks. RNA–X achieves the highest AUROC and AUPRC scores compared to existing RNA foundation models. RNA-X wo-head bars show the model's representations without representation head which consistently have lower performances.

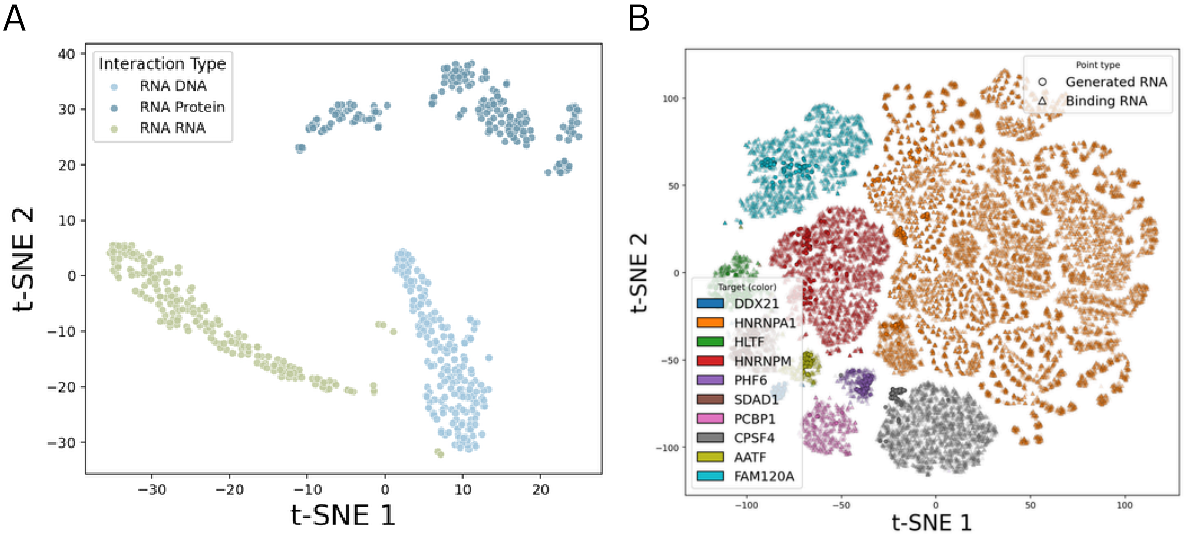


Supplementary Figure 8: **A** Visualization of RNA embeddings using t-SNE across three interaction types: RNA–protein, RNA–RNA, and RNA–DNA. The embeddings form distinct clusters by interaction type, indicating that RNA–X captures interaction-specific features. **B** Within RNA–protein interactions, embeddings of RNAs targeting the same protein cluster together, showing that the learned representations also encode fine-grained, target-specific information. Generated RNAs (shown in lighter shades) overlap closely with natural binding RNAs, which shows that RNA–X generates structurally and functionally coherent sequences in the learned embedding space.

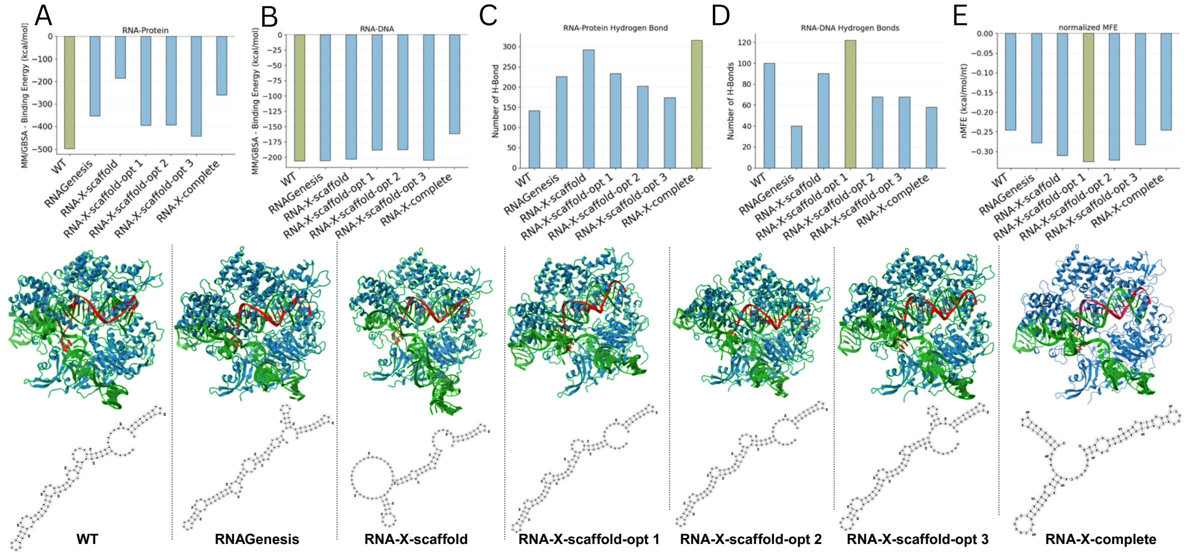


Supplementary Figure 9: **Multi-target RNA design with RNA–X in the Cas9–DNA–RNA complex.** MM/GBSA binding energies at the RNA–protein (**A**) and RNA–DNA (**B**) interfaces, hydrogen-bond counts at each interface (**C**, **D**), and the normalized MFE of the complex (**E**). The scaffold-optimized designs best preserve dual binding relative to the wild type (WT), while the *de novo* complete design stabilizes both interfaces less; all designs reach more negative normalized MFE than WT.

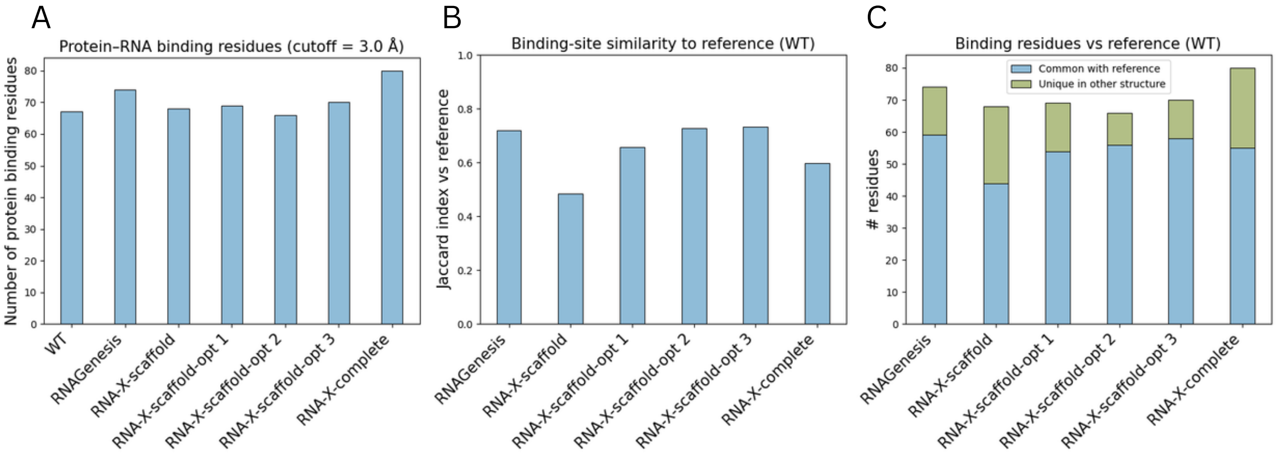


Supplementary Figure 10: **Structural comparison of WT and designed Cas9–DNA–RNA complexes.** **(A)** Number of protein residues that contact the RNA (3.0 Å cutoff). RNA-X-complete shows the largest number of contacts, while scaffold-optimized designs recover many WT-like interactions. **(B)** Binding-site similarity to the WT interface measured by the Jaccard index. Scaffold-optimized designs are most similar to the WT, whereas RNA-X-scaffold and RNA-X-complete share fewer WT contact residues. **(C)** Comparison of binding residues for each structure. Blue bars show residues shared with the WT interface, and green bars show residues unique to the designed complex. Scaffold-optimized designs preserve most WT contacts, while RNA-X-complete introduces many new interactions.

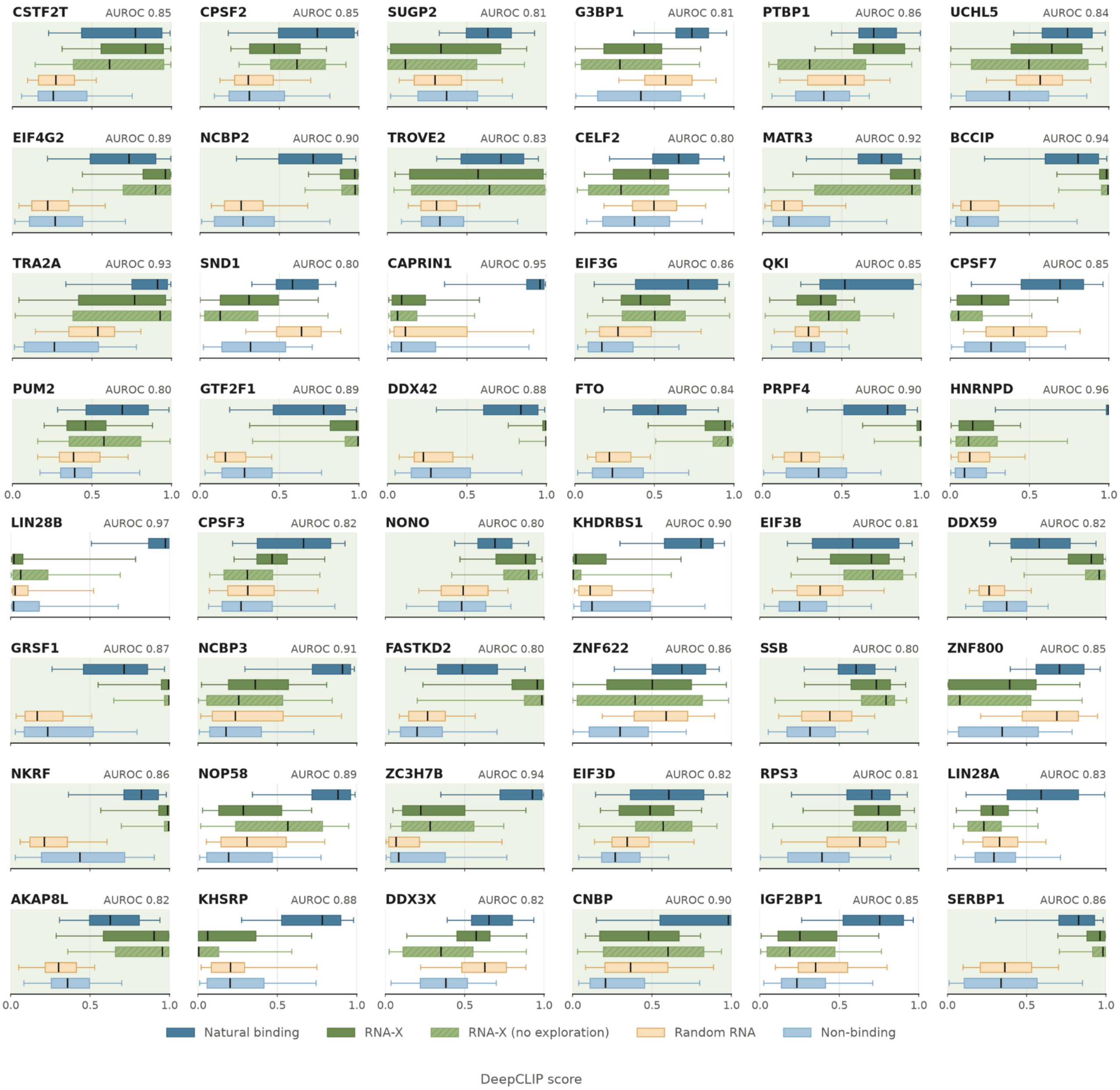


Supplementary Figure 11: **RNA design for 71 additional protein targets (part 1 of 2).** RNA–X was used to design binding RNAs for 71 proteins beyond the benchmarked set; no competing method could be run on these targets. Each panel shows one protein, with predicted binding scores for RNA–X designs, the ablated variant generated by iterative unmasking alone (RNA–X *no exploration*), the protein's known natural binders, randomly selected natural RNAs, and validated non-binders. The AUROC reported next to each protein name is the performance of that protein's dedicated DeepCLIP evaluator on its own held-out test set, and indicates how reliably the scores in that panel distinguish binders from non-binders; only proteins whose evaluator reached an AUROC of at least 0.80 were included. Panels shaded in green are those on which RNA–X designs score above the random-RNA control (50 of 71 across both parts). Remaining targets are shown in Supplementary Figure 12.

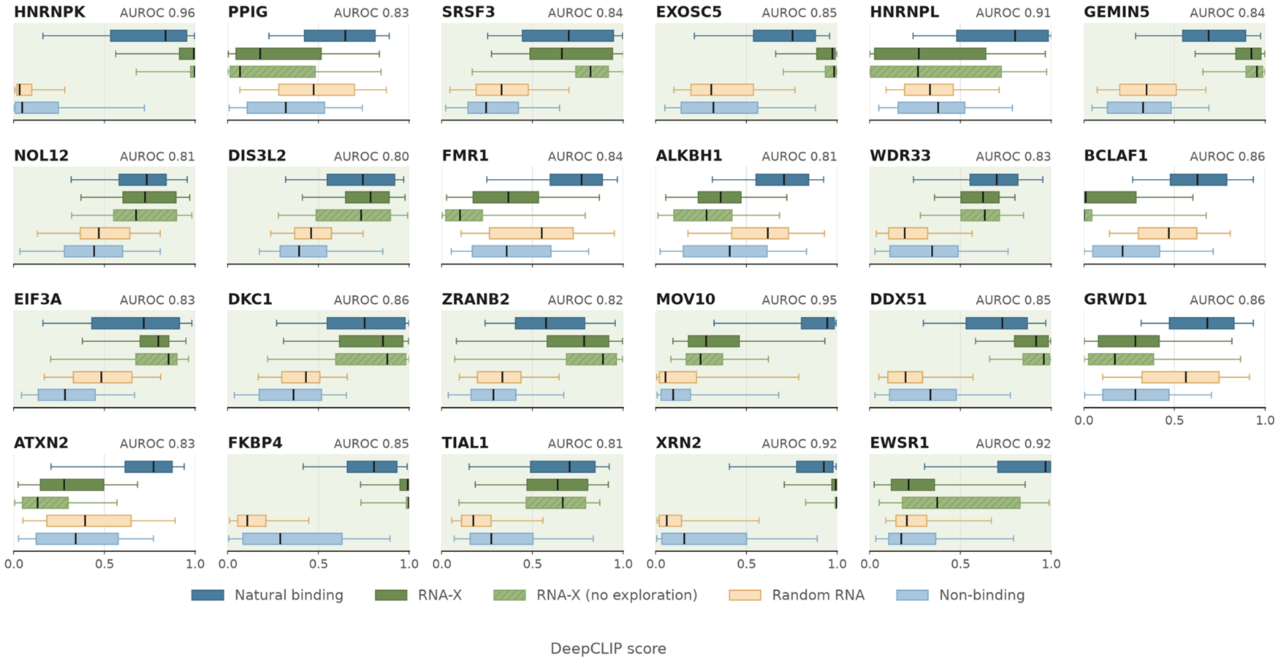


Supplementary Figure 12: **RNA design for 71 additional protein targets (part 2 of 2).** Continuation of Supplementary Figure 11, showing the remaining 23 protein targets. Panels, groups, AUROC annotation and shading are as defined there.

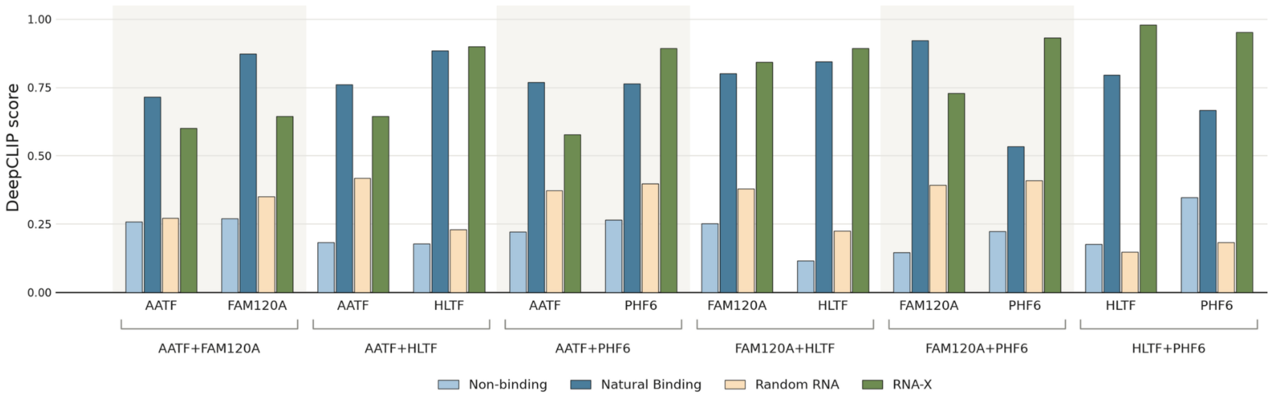


Supplementary Figure 13: **Multi-target RNA design for unseen protein pairs.** For each of the six pairwise combinations of the four proteins excluded from RNA–X's training data (AATF, FAM120A, HLTF, PHF6), a single RNA was designed conditioned on both target sequences simultaneously. Each design is scored independently with the DeepCLIP model of each protein in the pair and compared against that protein's known natural binders, randomly selected natural RNAs, and validated non-binders. Bars show mean predicted binding score.

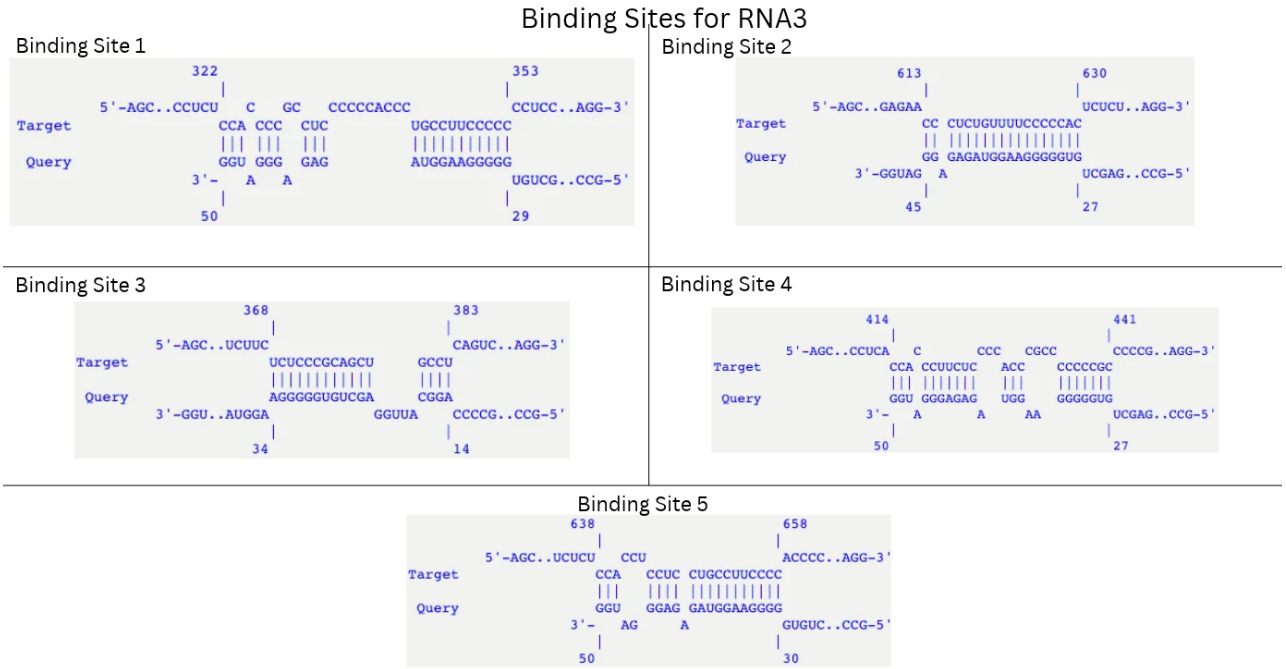


Supplementary Figure 14: **Predicted binding sites of RNA3 on the *AR* 5′UTR.** The five predicted interactions between RNA3 and the *AR* 5′UTR (Ensembl transcript AR-201), computed with IntaRNA. In each panel the target is shown $5' \rightarrow 3'$ (top) and the RNA3 query $3' \rightarrow 5'$ (bottom), with vertical bars marking base pairs and unpaired nucleotides shown offset. Numbers above and below each duplex give the first and last paired position on the target and the query, respectively. The five sites span positions 322–658 of the target and fall within the two accessible regions on which all four designs converge, and all five involve the 3′ half of RNA3.

### 3 Supplementary Tables

Supplementary Table 1: Summary of RNA-target interaction databases integrated in this study.

| **Type** | **Database** | **Total Interactions** | **Evidence** |
|---|---|---|---|
| RNA-Protein | NPInter v5.0[34] | 190,541 | Experimental |
| | RNAInter[16] | 33,338,687 | Computational |
| | RNAct[19] | 5.87 billion | Computational |
| | LncTarD[33] | 332 | Experimental |
| RNA-DNA | NPInter v5.0 | 4,703,412 | Experimental |
| | RNAInter | 69,368 | Computational |
| RNA-RNA | NPInter v5.0 | 1,832,429 | Experimental |
| | DIANA-LncBase v3[17] | 1,035,853 | Experimental |
| | StarBase v2[20] | 1,038 | Experimental |
| | LncTarD | 4,952 | Experimental |
| | miRDB[4] | 6,803,250 | Computational |

Supplementary Table 2: Summary statistics of filtered and clustered data for each interaction type.

| | **RNA-Protein** | **RNA-RNA** | **RNA-DNA** |
|---|---|---|---|
| Unique RNAs | 12,161,928 | 18,087 | 8,670 |
| Unique Targets | 76,794 | 103,806 | 2,986,307 |
| RNA Clusters | 3,035,451 | 10,095 | 8,371 |
| Target Clusters | 33,183 | 67,768 | 2,199,626 |
| Total Interactions | 95,877,665 | 3,052,842 | 3,115,730 |
| Experimental Interactions | 11,931,180 | 149,311 | 3,112,573 |
| Computational Interactions | 83,946,485 | 2,903,531 | 3,157 |

Supplementary Table 3: Non-redundant cluster interactions, redundancy, and overlap analysis.

| | **RNA-Protein** | **RNA-RNA** | **RNA-DNA** |
|---|---|---|---|
| Non-Redundant Cluster Interactions | 80,807,836 | 1,961,881 | 2,615,642 |
| Redundancy (Exp.) | 1.94 | 1.38 | 1.19 |
| Redundancy (Comp.) | 1.12 | 1.56 | 1.01 |
| Shared Cluster Interactions (Exp & Comp) | 293,077 | 2,520 | 5 |
| Exp.-only Cluster Interactions | 5,848,677 | 106,066 | 2,612,520 |
| Comp.-only Cluster Interactions | 74,666,082 | 1,853,295 | 3,117 |
| RNA Clusters (Exp & Comp) | 147,606 | 2,586 | 260 |
| Target Clusters (Exp & Comp) | 770 | 2,088 | 1,134 |

Supplementary Table 4: Number of sequences, clusters, and interactions in training and validation sets for each interaction type.

| | | **RNA-Protein** | **RNA-RNA** | **RNA-DNA** |
|---|---|---|---|---|
| Unique RNAs | Train | 11,267,967 | 17,933 | 8,655 |
| | Val | 1,017,627 | 10,252 | 6,684 |
| Unique Targets | Train | 72,941 | 98,520 | 2,835,872 |
| | Val | 3,853 | 5,218 | 150,435 |
| RNA Clusters | Train | 2,821,300 | 10,012 | 8,358 |
| | Val | 490,201 | 5,202 | 6,649 |
| Target Clusters | Train | 31,524 | 64,321 | 2,089,645 |
| | Val | 1,659 | 3,387 | 109,981 |
| Total Interactions | Train | 90,958,652 | 2,899,087 | 2,958,930 |
| | Val | 4,919,015 | 153,757 | 156,802 |
| Experimental Interactions | Train | 11,037,712 | 139,141 | 2,955,924 |
| | Val | 893,469 | 10,171 | 156,650 |
| Computational Interactions | Train | 79,920,940 | 2,759,946 | 3,006 |
| | Val | 4,025,546 | 143,586 | 152 |

Supplementary Table 5: **RNA sequences designed by RNA–X against the *AR* 5′UTR.** Transcript sequences are given 5′ → 3′. MFE and normalized MFE (nMFE) were computed with RNAfold at 37°C; $\Delta G_{\mathrm{hyb}}$ is the strongest predicted hybridization energy with the target 5′UTR computed with RNAup. Sites is the number of predicted binding sites on the target. The random sequence served as the negative control in all experiments.

| **Name** | **Transcript sequence** (5′ → 3′) | **Len.** | **GC** | **MFE** | **nMFE** | $\Delta G_{\mathrm{hyb}}$ |
|---|---|---|---|---|---|---|
| | | (nt) | (%) | (kcal/mol; nMFE per nt) | | |
| RNA1 | GGAAGGAAGCUCUGCCCCAGCAUCCCAGGUGGGGAGACAGUGGGAUGAGG | 50 | 64 | −14.40 | −0.288 | −21.50 |
| RNA2 | CCCUGCAGCCAGGGUGGUUGAGGCAGGGAUCUCUCUUUUCCUGGAAGGAG | 50 | 60 | −18.30 | −0.366 | −19.94 |
| RNA3 | GCCCUUCUGCCCCAGGCAUUGGAGCUGUGGGGGAAGGUAGAGAGGGAUGG | 50 | 64 | −18.50 | −0.370 | −18.50 |
| RNA4 | GAAGGAGUGGAGACCUCAGCUGGUUUGGGGCAUGAUGGAGGGCUUCCCCA | 50 | 60 | −11.50 | −0.230 | −18.07 |
| Random | CAUUCCCGGUACAGACUGAACAUUUUAACAAGGCGUUUUG | 40 | 40 | −2.10 | −0.053 | −8.60 |

Supplementary Table 6: **DNA oligonucleotides used for in vitro transcription.** The T7 promoter (TAATACGACTCACTATA) is underlined in the forward oligonucleotides and appears as its reverse complement at the 3′ end of the reverse oligonucleotides.

| **Oligo** | **Sequence** (5′ → 3′) |
|---|---|
| Random-F | TAATACGACTCACTATACATTCCCGGTACAGACTGAACATTTTAACAAGGCGTTTTG |
| Random-R | CAAAACGCCTTGTTAAAATGTTCAGTCTGTACCGGGAATGTATAGTGAGTCGTATTA |
| RNA1-F | TAATACGACTCACTATAGGAAGGAAGCTCTGCCCCAGCATCCCAGGTGGGGAGACAGTGGGATGAGG |
| RNA1-R | CCTCATCCCACTGTCTCCCCACCTGGGATGCTGGGGCAGAGCTTCCTTCCTATAGTGAGTCGTATTA |
| RNA2-F | TAATACGACTCACTATACCCTGCAGCCAGGGTGGTTGAGGCAGGGATCTCTCTTTTCCTGGAAGGAG |
| RNA2-R | CTCCTTCCAGGAAAAGAGAGATCCCTGCCTCAACCACCCTGGCTGCAGGGTATAGTGAGTCGTATTA |
| RNA3-F | TAATACGACTCACTATAGCCCTTCTGCCCCAGGCATTGGAGCTGTGGGGGAAGGTAGAGAGGGATGG |
| RNA3-R | CCATCCCTCTCTACCTTCCCCCACAGCTCCAATGCCTGGGGCAGAAGGGCTATAGTGAGTCGTATTA |
| RNA4-F | TAATACGACTCACTATAGAAGGAGTGGAGACCTCAGCTGGTTTGGGGCATGATGGAGGGCTTCCCCA |
| RNA4-R | TGGGGAAGCCCTCCATCATGCCCCAAACCAGCTGAGGTCTCCACTCCTTCTATAGTGAGTCGTATTA |

Supplementary Table 7: **qPCR primers used in this study.**

| **Primer** | **Sequence** (5′ → 3′) |
|---|---|
| HsARFL_qPCR_F | CAGCCTATTGCGAGAGAGCTG |
| HsARFL_qPCR_R | GAAAGGATCTTGGGCACTTG |
| HsKLK3_qPCR_F | CCACTGCATCAGGAACAAAAGC |
| HsKLK3_qPCR_R | ACCTGAAATACCTGGCCTGTG |
| HsActin_qPCR_F | GCAAAGACCTGTACGCCAAC |
| HsActin_qPCR_R | AGTACTTGCGCTCAGGAGGA |

Supplementary Table 8: **Reagents, kits and antibodies used in this study.**

| **Reagent** | **Source** | **Identifier** |
|---|---|---|
| RPMI-1640 | Gibco | 21875-034 |
| Fetal Bovine Serum, Tetracycline Free | Serana | S-FBF-SA-085 |
| Trypsin/EDTA, 0.25%, in HBSS with Phenol Red | Capricorn | TRY-3B |
| Lipofectamine™ MessengerMAX™ | Invitrogen | LMRNA001 |
| Penicillin/Streptomycin (100×) | Capricorn | CP25-8510 |
| Dulbecco's PBS, without Ca and Mg | Gibco | 14190-094 |
| TRUEzol | EcoTech | TR100 |
| Chloroform | Merck | 1024452500 |
| DEPC | Bioshop | DEP001 |
| 2× SYBR Green qPCR master mix | GLPBIO | GK10002 |
| TRIS Base | EcoTech | TBS10 |
| Sodium Dodecyl Sulfate (SDS), pellet | BIOMATIK | A3903-1KG |
| Acrylamide/Bis-Acrylamide (37.5:1), 30% (w/v) | BioBasic | S904R0K |
| Glycine | Bioshop | GLN001.1 |
| Methanol $\geq$ 99.7%, Ph Eur | ISOLAB | 947.043.2500 |
| Ethanol Absolute $\geq$ 99.9% | ISOLAB | LB.IS.920.026.2500 |
| 2-Propanol, Ph Eur, BP, JP, USP | ISOLAB | 961.023.2500 |
| Skim Milk Powder | Bioshop | SKI400.500 |
| Whatman GB003 gel blot paper | Cytiva | 10427818 |
| OPTI-MEM | Gibco | 31985070 |
| TEMED | AppliChem | A1148,0500 |
| Tween-20 | MP Biomedicals | 103168 |
| cOmplete Mini, EDTA-free | Roche | 78542300 |
| HiScribe® T7 High Yield RNA Synthesis Kit | New England BioLabs | E2040S/L |
| Supported Nitrocellulose membrane, 0.45 µm | GVS | 1212590 |
| Ultra-High Sensitivity ECL kit | GLPBIO | GK10008-200 |
| Bovine Serum Albumin | Sigma-Aldrich | A7030-10G |
| Bradford Reagent | EcoTech | BR05 |
| DTT | Molekula | 19733320 |
| EDTA Disodium | Multicell | 625-060-LG |
| Deoxycholic Acid Sodium Salt | BioBasic | DD0150 |
| NP-40 | AppliChem | A1694 |
| Sodium Chloride | ISOLAB | 969.033 |
| Protein Marker | EcoTech | PM12 |
| High-Capacity cDNA Reverse Transcription Kit | Thermo Scientific™ | 01206392 |
| AR (441), monoclonal, mouse | Santa Cruz Biotechnology | SC-7305 |
| $\beta$-actin, monoclonal, mouse | Merck (Sigma-Aldrich) | A1978-200UL |
| GOXMO HRP Affinity, Anti-Mouse Secondary Antibody | Invitrogen | A16066 |
